# Viral gene drive spread during herpes simplex virus 1 infection in mice

**DOI:** 10.1101/2023.12.07.570711

**Authors:** Marius Walter, Anoria K Haick, Rebeccah Riley, Paola A Massa, Daniel E Strongin, Lindsay M Klouser, Michelle A Loprieno, Laurence Stensland, Tracy K Santo, Martine Aubert, Matthew P Taylor, Keith R Jerome, Eric Verdin

## Abstract

Gene drives are genetic modifications designed to propagate efficiently through a population. Most applications rely on homologous recombination during sexual reproduction in diploid organisms such as insects, but we recently developed a gene drive in herpesviruses that relies on co-infection of cells by wild-type and engineered viruses. Here, we developed a viral gene drive against human herpes simplex virus 1 (HSV-1) and showed that it propagated efficiently *in vitro* and during HSV-1 infection in mice. We observed high levels of co-infection and gene drive-mediated recombination in neuronal tissues during herpes encephalitis as the infection progressed from the site of inoculation to the peripheral and central nervous systems. In addition, we found evidence that a superinfecting gene drive virus could recombine with wild-type viruses during latent infection. These findings indicated that HSV-1 achieves high rates of co-infection and recombination during viral infection, a phenomenon that is currently underappreciated. Overall, this study showed that a viral gene drive could spread *in vivo* during HSV-1 infection, paving the way toward therapeutic applications.

## Introduction

Herpesviruses are ubiquitous pathogens that establish lifelong infections and persistently infect most of the human population. Their large dsDNA genomes (100-200 kb) replicate in the nucleus and encode hundreds of genes. After primary infection, herpesviruses enter latency and occasionally reactivate, causing recurrent disease. Herpes simplex viruses (HSV) 1 and 2 first infect mucosal surfaces before spreading to the nervous system via axons. The viral genome remains latent in neurons in sensory and autonomic ganglia. Periodic reactivation causes lesions in the facial or genital area, which can be highly painful and stigmatizing. In addition, herpes encephalitis can be life-threatening in newborns and immunocompromised individuals (1, 2), and HSV-2 is a major risk factor for HIV acquisition (3). Antiviral drugs like acyclovir can block viral replication and reduce symptoms, but cannot eradicate th latent reservoir. HSV-1 and 2 lack vaccines or curative treatments and new therapeutic strategies for HSV diseases are critically needed.

During an infection, cells are often co-infected by several virions and therapeutic approaches that rely on viral co-infection have great potential. These viral interference strategies rely on defective viral particles that interfere with the replication of wild-type viruses after co-infection, thus reducing viral levels (4). Most viral interference approaches have focused on RNA viruses such as SARS-CoV-2, RSV, Influenza, Zika, Chikungunya, or HIV (5–10). We recently developed a system of viral interference against herpesviruses, using a technique of genetic engineering known as a gene drive (11, 12). Gene drives are genetic modifications designed to spread efficiently through a population (13, 14). They usually rely on CRISPR-mediated homologous recombination during sexual reproduction to efficiently spread a transgene in a population, and applications have been developed in insects to eradicate diseases such as malaria (13, 14). By contrast, the viral gene drive that we invented spread by the co-infection of cells with wild-type and engineered viruses (Fig. 1A). Specifically, upon co-infection with a gene drive virus expressing CRISPR-Cas9, the wild-type genome is cut and repaired by homologous recombination, producing new recombinant gene drive viruses that progressively replace the wild-type population. In a proof-of-concept with human cytomegalovirus (hCMV), we previously showed that a viral gene drive propagated efficiently *in vitro* in a population of unmodified viruses (11). Importantly, an attenuated gene drive virus could spread efficiently until the wild-type population had been eliminated, ultimately reducing viral levels. This represented a new way to engineer herpesviruses for research or therapeutic purposes.

**Figure 1:**
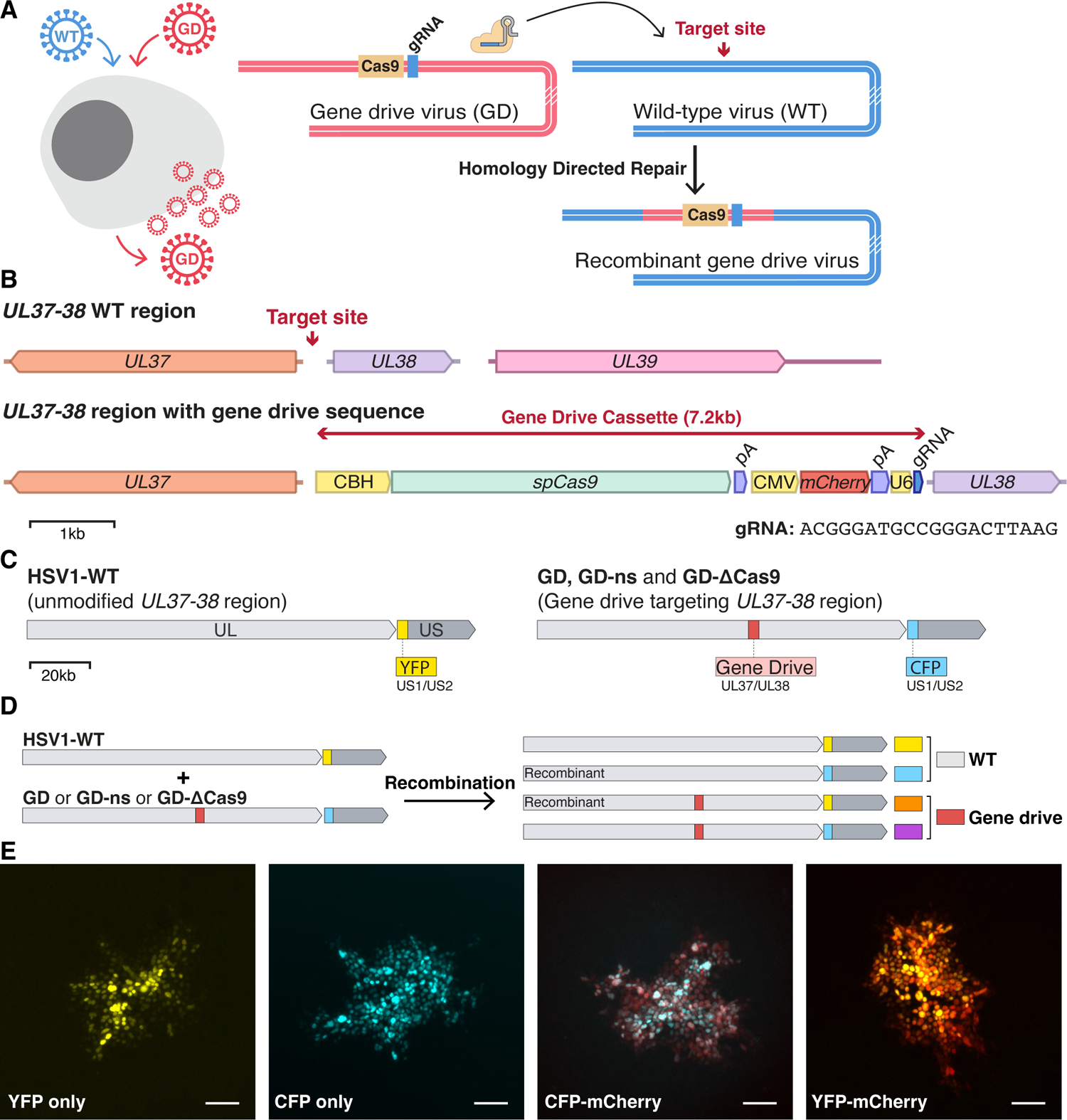
Design of a viral gene drive targeting HSV-1 *UL37-38* region. **A**. Gene drive viruses carry *Cas9* and a gRNA targeting the same location in a wild-type genome. After coinfection of cells by wild-type (WT) and gene drive (GD) viruses, Cas9 cleaves the wild-type sequence and homology-directed repair - using the gene drive sequence as a repair template-causes the conversion of the wild-type locus into a new gene drive sequence and the formation of new recombinant gene drive viruses. **B**. Modified and unmodified *UL37-38* region. The gene drive cassette was inserted between the *UL37* and *UL38* viral genes, and was composed of *spCas9* under the control of a CBH promoter followed by the SV40 polyA signal, a CMV promoter driving an mCherry reporter, followed by the beta-globin polyA signal, and a U6-driven gRNA. **C**. Localizations of the gene drive sequence and YFP/CFP reporters on HSV-1 genomes. GD represents a functional gene drive virus, GD-ns carries a non-specific gRNA, and Cas9 is deleted in GD-i1Cas9. UL/US: unique long/short genome segments. **D, E**. Recombination products and examples of viral plaques after cellular co-infection with HSV1-WT expressing YFP and gene drive viruses expressing mCherry and CFP.

The spread of a viral gene drive relies on cellular co-infection events, which occur at a high frequency in cell culture experiments. While numerical simulations suggest that a viral gene drive could still spread with low co-infection rates (11), the frequency of co-infection events during animal or human infections is poorly characterized. In mice, high levels of recombination are observed during herpes simplex encephalitis (15, 16), and attenuated viruses - which individually are harmless-can complement each other and cause severe disease (17, 18). Moreover, HSV strains circulating in humans exhibit evidence of extensive recombination (19–22). These indirect observations suggest that cells are often co-infected by several virions during HSV-1 infection. However, whether a viral gene drive could propagate *in vivo* remains unknown.

In the present study, we tested if a viral gene drive could spread in animal models of herpesvirus infection. HSV-1 is an important human pathogen with established infection protocols in mice, and we developed a gene drive that targeted a neutral region in the HSV-1 genome. This design was not intended to impact viral fitness or limit viral spread, but to investigate the potential of the technology *in vivo*. Our results show that a gene drive could spread efficiently during acute HSV-1 infection, particularly in neuronal tissues. Using fluorescently labeled viruses, we directly observed high levels of cellular co-infection in the brain and trigeminal ganglia. Finally, we show evidence that a superinfecting gene drive virus could recombine with wild-type genomes during latent infection, paving the way toward therapeutic applications.

## Results

### Design of a gene drive against HSV-1

We aimed to build a gene drive that would not affect viral infectivity and could spread efficiently into the wild-type population. We designed a gene drive targeting an intergenic sequence between the *UL37* and *UL38* genes, a region known to tolerate the insertion of transgenes with little or no impact on viral replication *in vitro* and *in vivo* (23, 24). We created a donor plasmid containing homology arms, *Cas9* (from Streptococcus pyogenes) under the CBH promoter, an *mCherry* fluorescent reporter under the CMV promoter, and a U6-driven gRNA targeting the intergenic *UL37*-*UL38* region (Fig. 1B). Importantly, to prevent self-cleaving, introduction of the gene drive cassette removed the gRNA target sequence from the construct. To create the gene drive virus (GD), Vero cells were co-transfected with purified HSV-1 viral DNA and the gene drive donor plasmid, and mCherry-expressing viruses created by homologous recombination were isolated by three rounds of plaque purification until a pure population was obtained. To follow recombination events between viral genomes, the gene drive virus also carried a cyan fluorescent reporter (CFP) inserted into another neutral region between the *US1* and *US2* viral genes (15, 25) (Fig. 1C). Similarly, we built two control viruses with non-functional CRISPR systems, one with a non-specific gRNA (GD-ns) that did not target HSV-1, and one without *Cas9* (GD-i1Cas9). Finally, to be used as a wild-type virus with an unmodified *UL37-UL38* region, we generated a virus expressing a yellow fluorescent reporter (YFP) from the same *US1-US2* region, hereafter referred to as HSV1-WT or simply WT (Fig. 1C). All the viruses described here originated from the highly neurovirulent HSV-1 strain 17+.

Recombination between gene drive and wild-type genomes can result in four different genome configurations expressing the different fluorescent reporters, which can be followed by plaque assay (Fig. 1D and 1E). In highly susceptible cell lines such as Vero cells, viral plaques originate from single virions, which allows reconstituting the recombination history of individual viral genomes. Plaques expressing YFP only, or CFP and mCherry together, represent the original WT and gene drive viruses, respectively, while plaques expressing CFP only, or YFP and mCherry together, represent recombination products that exchanged the gene drive cassette.

### Gene drive spread in cell culture

First, we tested if our gene drive could spread *in vitro* and conducted co-infection experiments in cell culture. Some cell lines, in particular Vero cells or other epithelial cells, efficiently restrict co-infection by a mechanism known as superinfection exclusion (26, 27). However, we found that mouse neuronal N2a cells could sustain high levels of co-infection (Supplementary Fig. S1). Thus, co-infection experiments were conducted in N2a cells, while plaque assays were performed in Vero cells. Importantly, the WT, GD, GD-ns and GD-i1Cas9 viruses individually replicated with similar dynamics in N2a cells, showing that insertion of the gene drive cassette in the *UL37-UL38* region did not affect infectivity *in vitro,* as expected (Fig. 2A).

**Figure 2:**
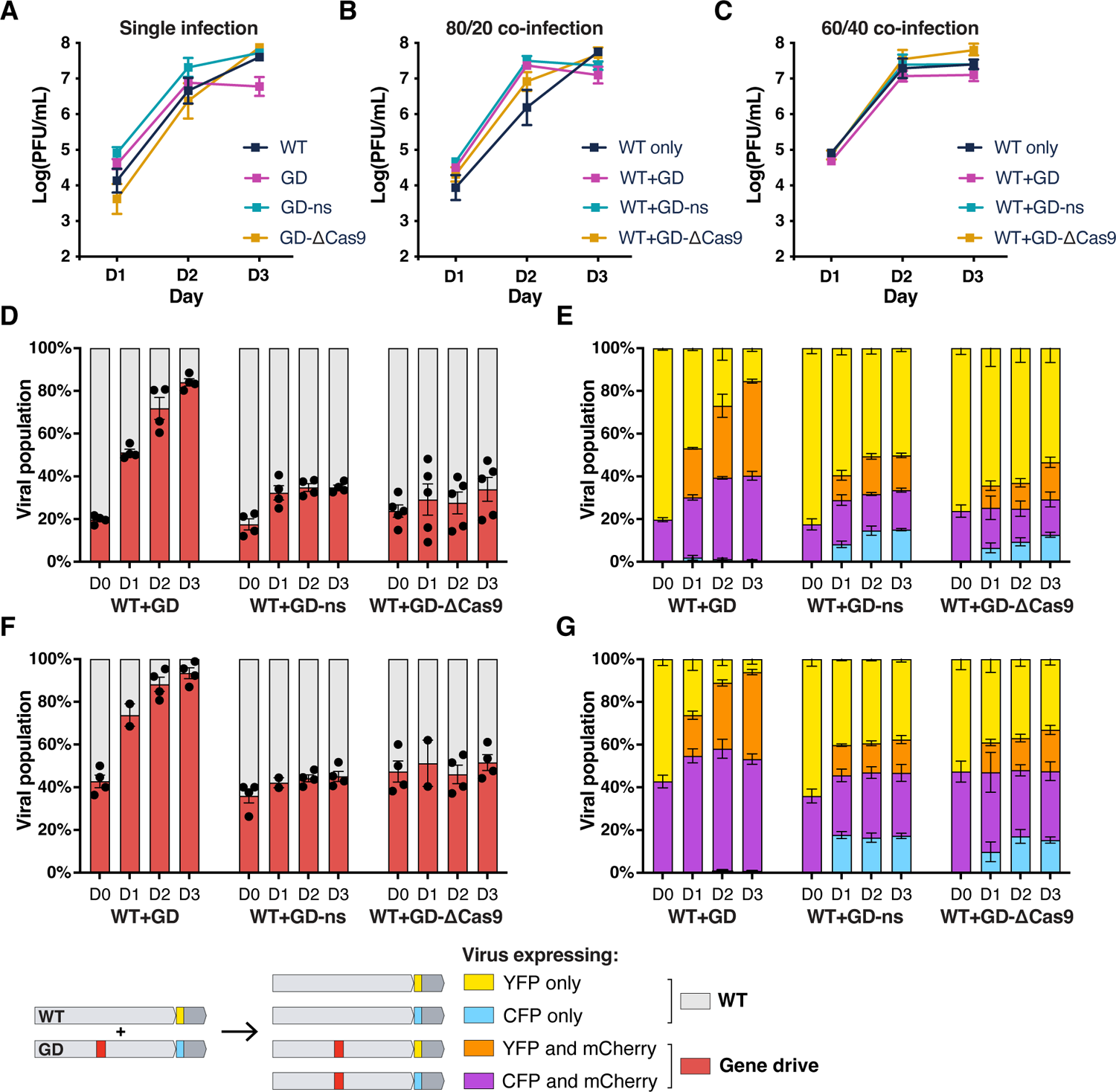
Gene drive spread in cell culture. **A**. Viral titers in the supernatant after infection of N2a cells with WT, GD, GD-ns or GD-i1Cas9. Cells were infected with a single virus at MOI=1. n=4. **B, C**. Viral titers in the supernatant after co-infection of N2a cells with WT+GD, WT+GD-ns or WT+GD-i1Cas9, with a starting proportion of gene drive virus of 20% (B) or 40% (C). MOI=1, n=4. **D-G**. Evolution of the viral population after co-infection with WT+GD, WT+GD-ns or WT+GD-i1Cas9, with a starting proportion of gene drive virus of 20% (D, E) or 40% (F, G). Panels D and F show the proportion of viruses expressing mCherry, representing gene drive virus. Panels E and G show the proportion of viruses expressing the different fluorophore combinations. Viral titers are expressed in log-transformed PFU (plaque-forming unit) per mL of supernatant. Error bars represent the standard error of the mean (SEM) between biological replicates. n=4. Source data are provided as a Source Data file.

To test if the gene drive could efficiently spread in the viral population, N2a cells were co-infected with WT+GD, WT+GD-ns, or WT+GD-i1Cas9. Cells were infected at a combined MOI of 1 with an initial proportion of gene drive virus of 20% (Fig. 2B, 2D and 2E). Titers and proportion of progeny viruses expressing the different fluorescent markers were measured by plaque assay, from day one to day three post-infection. The proportion of viruses expressing mCherry, which represents gene drive viruses, increased from 20% to 85% when cells were co-infected with WT+GD (Fig. 2D). Importantly, YFP-only viruses disappeared and were replaced by recombinant viruses expressing both YFP and mCherry, representing 45% of the final population (Fig. 2E). This indicated that the WT population had been converted to new recombinant gene drive viruses, as anticipated. The proportion of viruses expressing both CFP and mCherry, which represent the original gene drive virus, increased slightly from 20% to 40% (Fig. 2E). By contrast, in the control experiments with GD-ns or GD-i1Cas9, the proportion of mCherry-expressing viruses did not change and remained close to its initial value of 20% (Fig. 2D). In these control experiments, around 10% of viruses expressed both YFP and mCherry at day 3, but this population of recombinant viruses was mirrored by viruses that had lost mCherry and expressed CFP only (Fig. 2E). These represented viruses that had exchanged their YFP and CFP regions in a nonspecific manner. Importantly, CFP-only viruses were not observed after co-infection with WT+GD, highlighting that the efficient incorporation of the gene drive cassette into unmodified genomes is a unilateral and targeted process requiring both Cas9 and a specific gRNA.

To confirm these observations, we repeated these co-infection experiments with an initial proportion of gene drive virus of 40% (Fig. 2C, 2F, and 2G). With this higher starting point, the gene drive achieved almost complete penetrance and the proportion of mCherry-expressing viruses reached 95% after 3 days, with the population of wild-type viruses expressing YFP-only being converted to recombinant gene drive viruses expressing both YFP and mCherry. As observed above, In the WT+GD-ns or WT+GD-i1Cas9 control experiments, the proportion of mCherry-expressing viruses remained constant at around 40%, and approximately 15% of YFP+mCherry and CFP-only viruses symmetrically appeared by CRISPR-independent recombination. This further showed that a gene drive could spread efficiently in the wild-type population and that the drive was mediated by a functional CRISPR system.

Together, these results indicated that a gene drive could be designed against HSV-1 and spread efficiently *in vitro.* Importantly, both during single and co-infections, viral growth of WT and GD viruses followed similar dynamics and the proportion of viruses expressing CFP remained constant (Fig. 2). Thus, the rapid increase of recombinant viruses expressing both mCherry and YFP could not be explained by a higher fitness of the GD virus, but resulted from efficient CRISPR-directed homologous recombination. These results align with the findings of a parallel study (28) and expanded on our previous work with hCMV, showing that a viral gene drive could be developed in a second, unrelated, herpesvirus.

### Gene drive spread during herpes encephalitis

Next, we tested if the gene drive could spread during acute HSV-1 infection in mice. Previous studies showed that HSV-1 naturally sustains high levels of recombination in the mouse brain during herpes simplex encephalitis (15). Thus, we hypothesized that a gene drive could spread efficiently in this context. We used a well-established infection model, where HSV-1 is inoculated intravitreally in the eye and infects the retina and other ocular tissues before propagating to the nervous system via cranial nerves (Fig. 3A). Specifically, HSV-1 travels to the brain via the optic, oculomotor and trigeminal nerves (cranial nerve CN II, III, and V respectively), either directly or indirectly by first infecting ganglionic neurons of the peripheral nervous system. In particular, HSV-1 infects the trigeminal ganglia (TG) before reaching the brain stem. In the following experiments, a total of 10^6^ plaque-forming units (PFU) were inoculated intravitreally in the left eye, and tissues were collected, dissociated, and analyzed by plaque assay after four days. Five to eight-week-old male and female Balb/c mice were used and we did not observe any differences between sexes. Importantly, individual infections with WT, GD or GD-ns reached similar titers in the eye, the TG and the brain, showing that the gene drive cassette in the *UL37-UL38* region did not impact infectivity *in vivo* (Fig. 3B).

**Figure 3:**
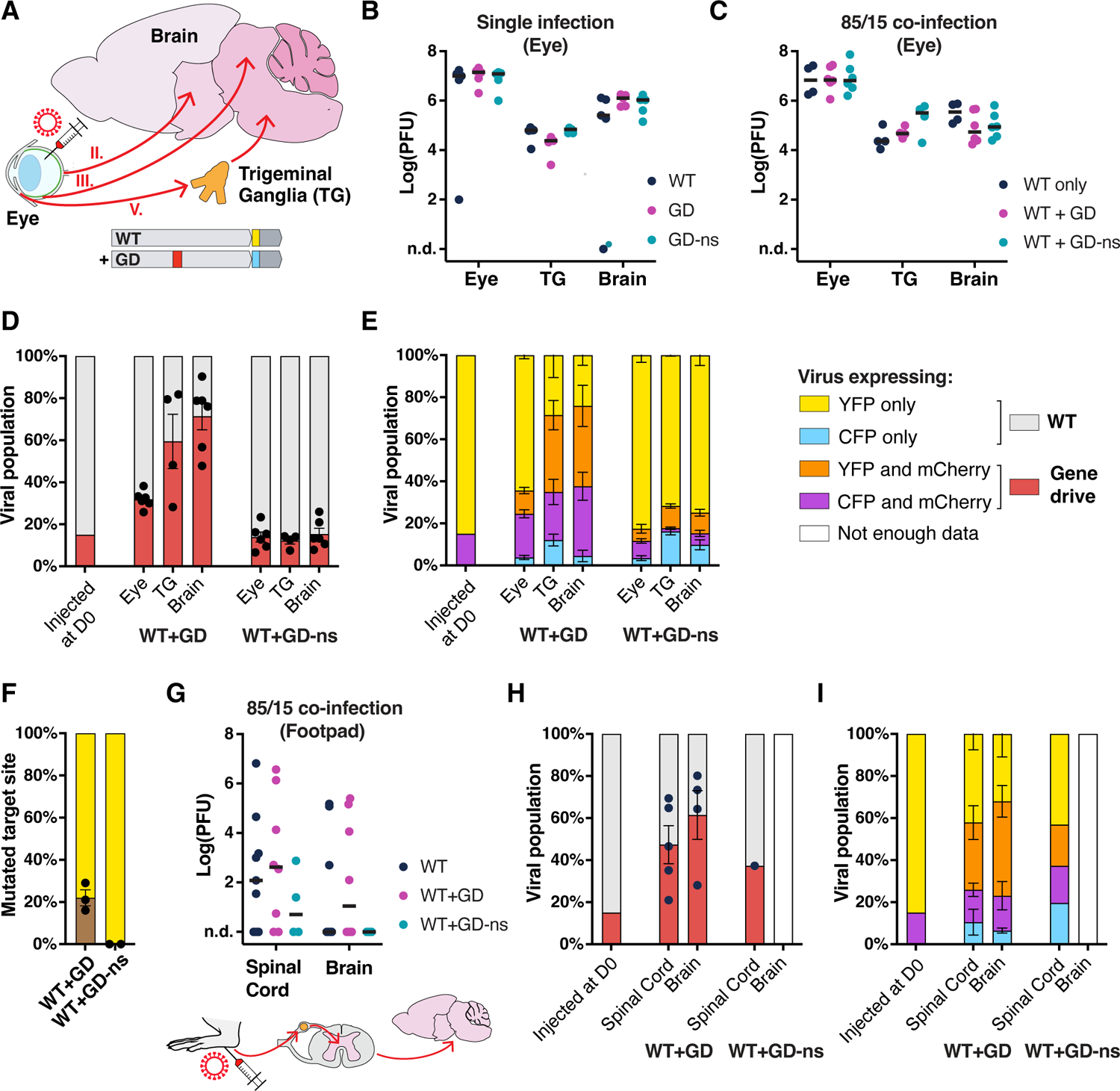
Gene drive spread during herpes simplex encephalitis. **A**. Infection routes along optic, oculomotor and trigeminal nerves (cranial nerves II, III and V, respectively) following ocular inoculation of HSV-1. Male and female Balb/c mice were infected with 10^6^ PFU in the left eye. **B, C**. Viral titers after four days in the eye, TG and whole brain after (B) infection with a single virus, n=5, or (C) with a starting proportion of gene drive virus of 15%, n=6. **D, E**. Viral population in the eye, TG and whole brain after co-infection with WT+GD or WT+GD-ns, after four days. n=6. **F**. Proportion of viral genomes with a mutated target site in the brain after four days. n=3. **G**. Viral titers in the spinal cord and brain after inoculation of WT, WT+GD, or WT+GD-ns in the right hind leg footpad, after 5-7 days. n=8 for WT and WT+GD, n=4 for WT+GD-ns. **H, I**. Viral population in the spinal cord and whole brain after co-infection with WT+GD or WT+GD-ns, after 5-7 days. n=5 for WT+GD, n=1 for WT+GD-ns. Viral titers are expressed in log-transformed PFU. In panels B, C, G, black lines indicate the median. n.d.: non-detected. Panels D, E, F, H and I show the average and SEM between biological replicates. Source data are provided as a Source Data file.

To test if the gene drive could efficiently spread *in vivo*, mice were inoculated intravitreally with WT only, WT+GD, or WT+GD-ns, with an initial proportion of gene drive virus of 15% (Fig. 3C-3E). Total viral titers in the eye, TG and brain after four days were indistinguishable between the different conditions, showing that the gene drive did not perturb the overall dynamics of infection (Fig. 3C). The population of gene drive viruses expressing mCherry increased from 15% to 30% in the eye, to 60% in the TG, and 70% in the brain, respectively.

We observed a high variation between replicates, with the percentage of gene drive viruses in the brain ranging from 50% to 90%, with a median of 77% (Fig. 3D). Furthermore, and as observed *in vitro,* wild-type viruses expressing YFP-only were converted to recombinant gene drive viruses expressing YFP and mCherry, representing 40% of the final population. The proportion of original gene drive viruses expressing CFP and mCherry increased slightly, from 15% to 20% in the TG, and 30% in the brain, respectively (Fig. 3E). Critically, in the control experiment with GD-ns, the proportion of gene drive viruses did not change and remained close to its initial value around 15% in all tissues, with a similar proportion of YFP+mCherry and CFP-only viruses appearing by CRISPR-independent recombination (Fig. 3E). Altogether, this important result shows that the gene drive efficiently spread in the viral population as the infection progressed to the brain, with recombinant gene drive viruses increasing from 15% to 70% in four days.

Gene drive propagation relies on efficient homologous recombination after CRISPR cleavage, but DNA double-strand breaks can also be repaired by non-homologous end joining (NHEJ), resulting in small insertions and deletions that render viruses resistant to the drive (12, 28). After PCR and Sanger sequencing of infectious viruses isolated from the brain, we found that 20% of the remaining target sites had been mutated by NHEJ (Fig. 3F). Since the gene drive represented 70% of viruses at this point, this result suggested that drive-resistant viruses represented around 6% of the total viral population. This indicated that the drive did not achieve full penetrance after four days, and confirmed the observation made with hCMV that mutagenic repair by NHEJ is infrequent compared to homologous recombination during gene drive spread (11, 12).

We next tested if our gene drive could spread in another infection model, where HSV-1 is inoculated in the hind leg footpad. In this model, HSV-1 travels to the spinal cord via the sciatic nerve, and finally to the brain. 10^6^ PFU of WT, WT+GD or WT+GD-ns were inoculated in the right hind footpad, with an initial proportion of gene drive virus of 15%. Tissues were collected five to seven days later and analyzed by plaque assay (Fig. 3G-I). Around half the mice did not show any symptoms of infection and had no detectable virus in the spinal cord and the brain, while others developed severe neurological symptoms with very high titers in both tissues (Fig. 3G). In animals with detectable virus, the average proportion of gene drive viruses reached 60% in the brain, with a range between 30% and 80%, and 50% in the spinal cord (Fig. 3H). Once again, wild-type viruses expressing YFP-only had been converted to recombinant gene drive viruses expressing both YFP and mCherry, while the population of viruses expressing CFP and mCherry remained constant (Fig. 3I). Only one control animal infected with WT+GD-ns developed a detectable infection, with mCherry-expressing viruses reaching 35% in the spinal cord. These results confirmed that a gene drive could spread after inoculation via a second route of HSV-1 infection. Altogether, these findings showed that a gene drive can propagate efficiently *in vivo* during acute HSV-1 infection in mice, and that the drive was mediated by a functional CRISPR system.

### High heterogeneity during gene drive spread in the brain

HSV-1 travels to the brain via different neuronal pathways, and we hypothesized that following gene drive propagation in different regions of the brain could bring novel insight into the dynamics of HSV-1 infection and recombination. After intravitreal inoculation, HSV-1 infects retinal neurons and travels via the optic nerve (CN II) to the hypothalamus and the thalamus - both part of the interbrain-before reaching the cortex through visual pathways. Secondary branches of the optic nerve also connect to the midbrain - the rostral part of the brain stem. After infecting other ocular tissues and specifically the ciliary ganglion, HSV-1 separately reaches the midbrain via the oculomotor nerve (CN III). Finally, HSV-1 travels via the trigeminal nerve (CN V) to the TG and then to the brain stem (Fig. 4A).

**Figure 4:**
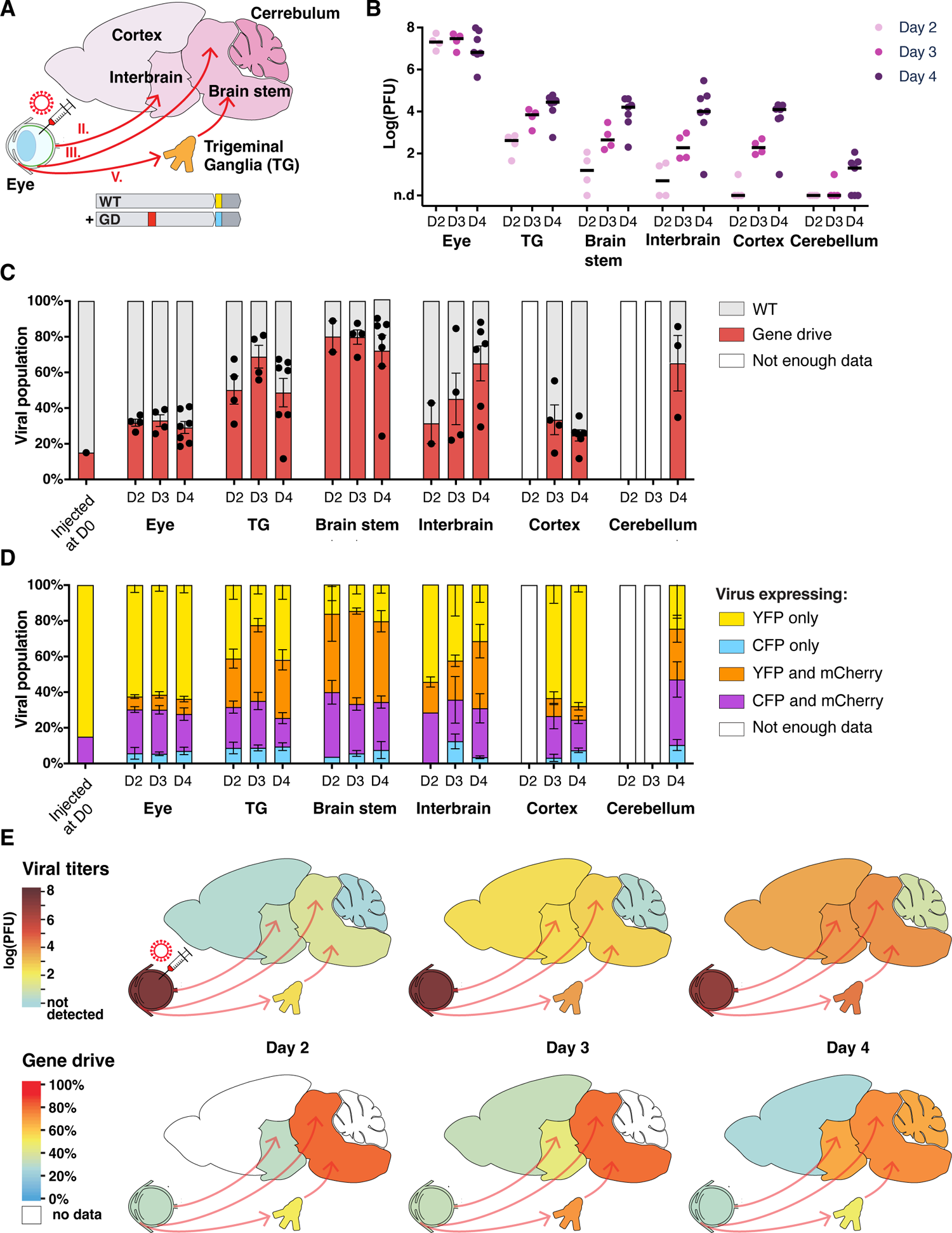
High heterogeneity between brain regions during gene drive spread. **A**. Infection routes following ocular inoculation of HSV-1. Male and female Balb/c mice were co-infected with 10^6^ PFU of WT+GD in the left eye, with a starting proportion of gene drive virus of 15%. **B**. Viral titers over time. Black lines indicate the median. n.d.: non-detected. **C, D**. Proportion of gene drive viruses over time. Data show the average and SEM between biological replicates. **E**. Heatmap summarizing panels B and C. n=4 for day 2 and 3, n=6 for day 4. Source data are provided as a Source Data file.

To investigate tissue-specific differences in gene drive propagation, Balb/c mice were inoculated intravitreally with 10^6^ PFU of WT+GD, with an initial proportion of gene drive virus of 15%. Viral titers and gene drive-directed recombination were measured by plaque assay in the eye, TG, brain stem, interbrain, cortex and cerebellum at days two to four post-infection (Fig. 4). Viral titers increased progressively throughout the brain, first reaching the interbrain and brain stem after 2 days and then spreading to the cortex and cerebellum (Fig. 4B and summary heatmap in Fig. 4E, upper panel). As described above, the proportion of gene drive viruses remained low in the eye and reached around 60% in the TG (Fig. 4C-4E). Interestingly, gene drive levels varied greatly between the different brain regions. The gene drive reached 80% in the brain stem after only two days, and progressively increased to 65% in the interbrain and cerebellum, with a penetrance of up to 90% in some animals. By contrast, gene drive levels remained low in the cortex, at 25% on average. As observed above, in the TG, brain stem and interbrain - but not in the eye and cortex, viruses expressing YFP-only had been converted to recombinant gene drive viruses expressing both YFP and mCherry (Fig. 4D). Together, this showed that gene drive spread in the brain was highly heterogeneous, with almost complete penetrance in some regions and barely any in adjacent ones.

This heterogeneity gives an interesting insight into the dynamic of HSV-1 infection. Gene drive propagation relies on cellular co-infection and one intuitive hypothesis is that higher viral levels would increase the probability that cells are co-infected by several virions, and, thus, that recombination levels should positively correlate with viral titers. However, no correlation between viral titers and recombination was observed (Supplementary Fig. S2). For example, HSV-1 levels were the highest in the eye, but gene drive levels remained the lowest in this organ. Similarly, HSV-1 reached similar titers in the brain stem, interbrain and cortex, but almost no recombination occurred in the cortex while high levels were observed in the brain stem and interbrain (Fig. 4D). This shows that recombination does not simply correlate with viral titers, as could be expected, but is likely explained by other tissue-specific cellular or viral mechanisms.

### High levels of cellular co-infection in the TG and the brain

The results described above suggest that cells are frequently co-infected by several virions during HSV-1 infection. Thus, we aimed to directly measure co-infection levels during HSV-1 infection, as this would provide important insight into the basic biology that supports gene drive propagation.

HSV-1 and related viruses expressing fluorescent proteins have long been used to probe neuronal pathways of the visual system (29–31). To investigate cellular co-infection, Balb/c mice were infected ocularly with equal amounts of three different viruses, expressing either YFP, CFP or RFP from the same *US1/US2* locus (Fig. 5A). The fluorescent reporters carried nuclear localization signals and were not incorporated into virions, and, thus, marked infected nuclei. Mice were injected intravitreally with a total of 10^6^ PFU in the left eye and dissected four days later. Fluorescence was observed directly on frozen sections without staining. Strikingly, we observed very high levels of co-infection in the TG (Fig. 5) and different regions of the brain (Fig. 6), with cells often co-expressing two or more fluorophores.

**Figure 5:**
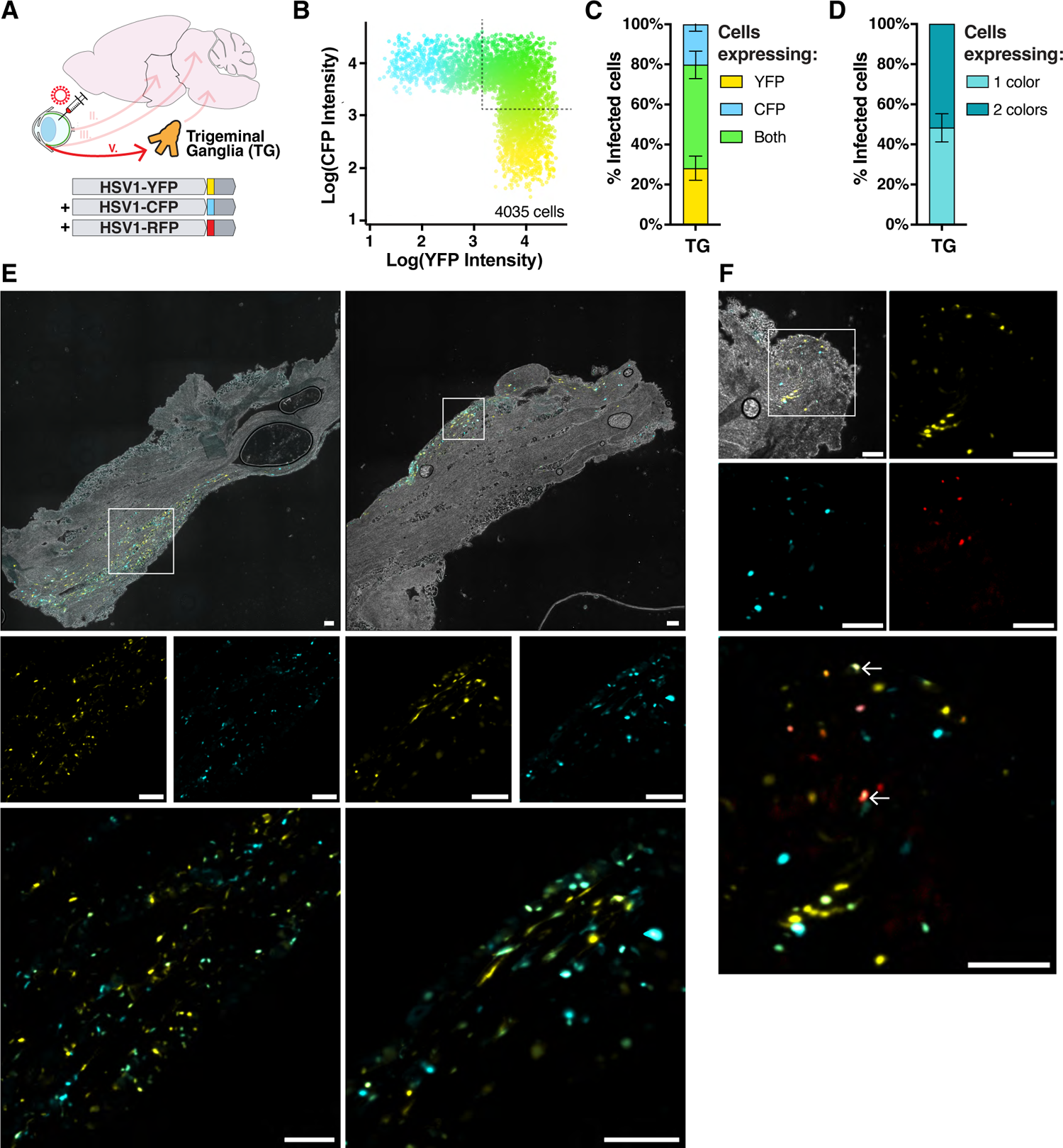
High levels of co-infection in the TG during HSV-1 infection. **A**. Balb/c mice were co-infected with equivalent amounts of three viruses expressing YFP, CFP and RFP, respectively, with a total of 10^6^ PFU in the left eye. **B**. YFP and CFP cellular intensity after machine learning-assisted cell segmentation of TG sections. Datapoints represent individual cells and were colored by converting YFP and CFP signals into the CYMK color space. 4035 cells were detected, originating from 53 images and n=4 animals. **C**. Percentage of infected cells expressing YFP, CFP, or both. n=4. **D**. Percentage of infected cells expressing one or two fluorescent markers. n=4. **E**, **F**. Representative images of TG sections, highlighting high levels of co-infection. Scale bars: 100 µm. Arrows indicate cells co-expressing YFP, CFP and RFP together.

**Figure 6:**
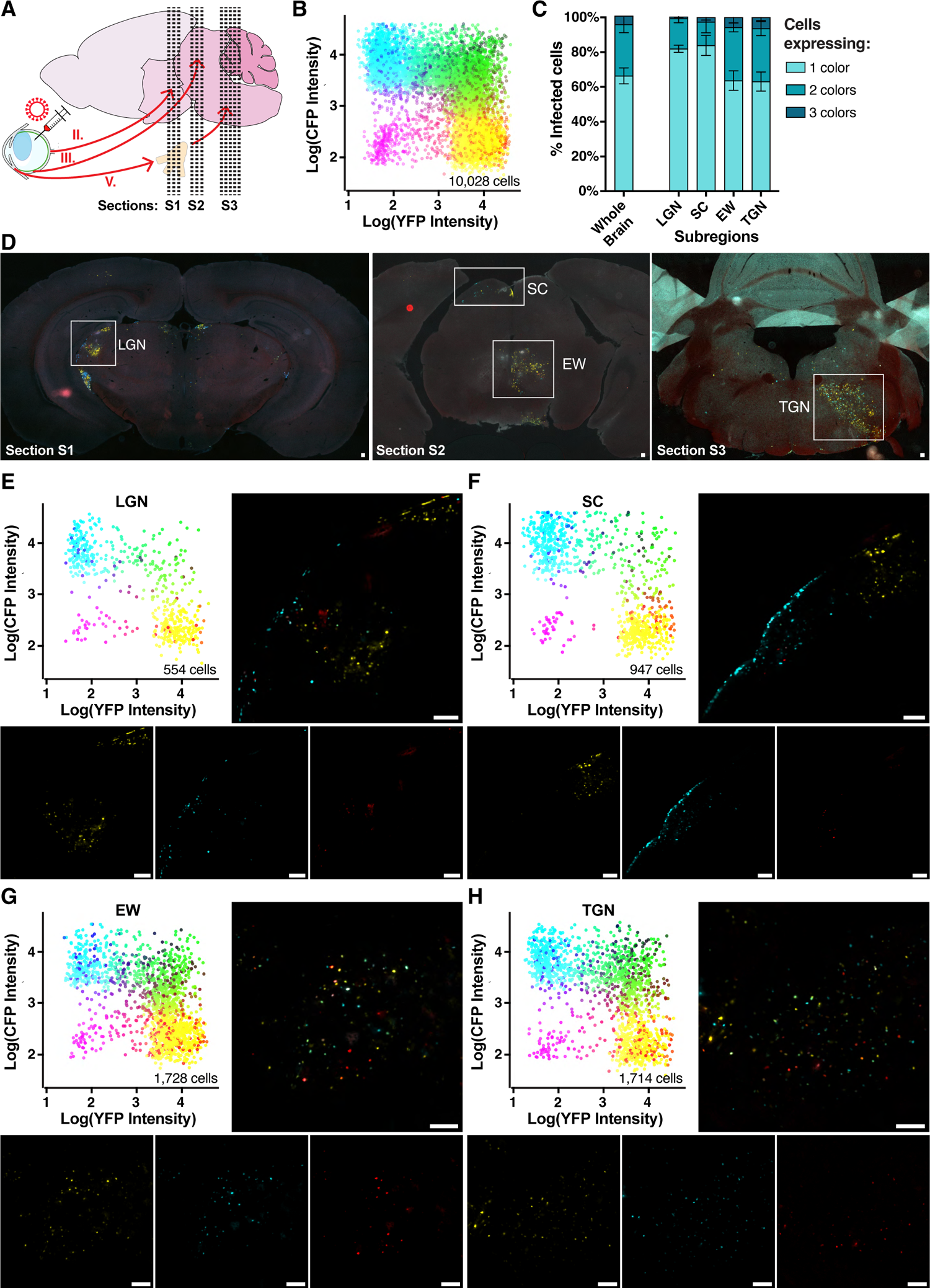
High levels of co-infection in the brain during HSV-1 infection. **A**. Images of brain sections were collected in three regions in the thalamus, midbrain and brain stem after ocular infection. **B**. YFP, CFP and RFP cellular intensity after machine learning-assisted cell segmentation of brain sections. Datapoints were colored by converting YFP, CFP and RFP signals into the CYMK color space. 10,028 cells were detected, originating from 95 images and n=3 animals. **C**. Percentage of infected cells expressing one, two, or three fluorescent markers, both in the whole brain and in specific subregions. n=3. **D**. Representative images of the brain in the thalamus (sections S1), midbrain (sections S2) and brain stem (section S3). **E**-**H**. Representative images and summary of co-infection patterns in specific subregions. LGN: lateral geniculate nucleus; SC: superior colliculus; EW: Edinger-Westphal nucleus; TGN: trigeminal nerve nuclei. Scale bars: 100 µm.

In the TG, YFP and CFP were expressed at high levels, but the RFP signal was weaker and rarely above background. Thus, for the TG, we restricted our analysis to YFP and CFP. We observed widespread expression distributed over the entire length of the TG on some sections, or more tightly localized clusters in others (Fig. 5E). Areas with the strongest signal likely localized to the neuron-rich ophthalmic division of the TG, but the widely disseminated expression indicated that HSV-1 had spread to other areas. Importantly, a majority of infected cells appeared to express both YFP and CFP, indicating extensive cellular co-infection in the TG (Fig. 5E). To quantify co-infection precisely, we used machine learning to automatically segment cells and measure YFP and CFP intensity on TG sections (Fig. 5B-5D, Supplementary Fig. S3). A total of 4035 cells was detected, with a clear separation between YFP and CFP signals and high consistency between replicates. We found that an average of 52% of cells expressed both YFP and CFP, with a range of 45% to 60% between replicates. Since RFP was excluded from this analysis, this was probably an underestimate of the co-infection frequency. In fact, in the few images with strong RFP signal, we detected numerous cells co-expressing RFP, YFP and/or CFP, with instances of cells expressing all three markers (Fig. 5F, Supplementary Fig. S3F). Together, this showed that more than 50% of cells were co-infected by two or more virions in the TG.

We then repeated this analysis in the brain (Fig. 6, Supplementary Fig. S4-S9). We detected strong fluorescence along well-characterized routes through the optic, oculomotor and trigeminal nerves (Fig. 6D, Supplementary Fig. S4). In particular, fluorescence was observed: 1) in the lateral geniculate nucleus (LGN) in the thalamus and the superior colliculus (SC) in the midbrain, where most axons of the optic nerves terminate; 2) in the Edinger-Westphal nucleus (EW), one of the two nuclei of the oculomotor nerve in the midbrain; and 3) throughout the hindbrain, likely corresponding to trigeminal nerve nuclei (TGN). Fluorescence was also detected in other areas associated with visual pathways such as the optic tract, olivary pretectal nucleus, suprachiasmatic nuclei or visual cortex (Supplementary Fig. S4). Additionally, disseminated fluorescence was detected over wide areas not easily associated with the visual system, with important variation between replicates. This likely corresponded to the secondary or tertiary spread of HSV-1 throughout the brain. Images were collected in three main regions in the thalamus, midbrain, and hindbrain (Fig. 6A). This time, RFP had a stronger signal and was included in the analysis. After machine learning-assisted segmentation, a total of 10,028 cells were analyzed over three brains. RFP was observed in 10-15% of infected cells, while YFP and CFP were detected in equivalent proportions in 40-60% of cells (Supplementary Fig. S5). When analyzing all images together, we observed a high level of co-infection. In total, 29% of cells expressed two colors, and 5% of cells had three colors, with results highly consistent between replicates (Fig. 6B, 6C, and Supplementary Fig. S5). This confirmed the high level of co-infection found in the TG, with more than 34% of cells co-infected by two or more virions in the whole brain.

We next focused our analysis on the brain regions associated with primary HSV-1 spread, namely the LGN, SC, EW and TGN. Interestingly, the LGN and SC, where the optic nerve terminates, had relatively low co-infection levels, around 20%. By contrast, the EW and TGN, where the oculomotor and trigeminal nerves terminate, respectively, had much higher co-infection levels, around 40% (Fig. 6C). This visually correlated with very different infection patterns. In the LGN and SC, well-separated and tight foci expressing only one color were observed, with co-infected cells at the boundaries (Fig. 6E, 6F, closeups and additional examples in Supplementary Fig. S6 and S7). This was reminiscent of viral plaques in cell culture and may suggest clonal spread from a single infected cell. By contrast, in the EW and TGN, infected cells were broadly disseminated, with few infected cells touching each other. The different colors were uniformly distributed and cells with one, two, or three colors were observed without evidence of spatial clustering (Fig. 6G, 6H, Supplementary Fig. S8 and S9). These differences in co-infection patterns are intriguing. Together with the high heterogeneity observed during gene drive spread (Fig. 4), these observations suggested that infection dynamics vary significantly depending on the viral propagation route. This will form the basis of future investigations.

Altogether, this analysis revealed that neuronal tissues sustain high levels of co-infection during HSV-1 infection, with more than 50% and 40% of cells infected with two or more viruses in the TG and specific brain regions, respectively.

### Gene drive spread during latent infection

Our results showed that a gene drive can spread efficiently during acute infection. Next, we investigated if a gene drive could also spread in the context of a latent infection, as this would open important avenues for therapeutic interventions. After primary oro-facial infection, HSV-1 typically establishes latency in the TG and other peripheral ganglia.

Following reactivation, HSV-1 travels back to the mucosal surface, causing lesions or shedding asymptomatically. In the following experiments, we used an ocular model of latent infection and drug-induced reactivation in mice to test if a gene drive virus - administered at a later time point, thus “superinfecting”-could target and recombine with latent HSV1-WT (Fig. 7 and 8).

**Figure 7:**
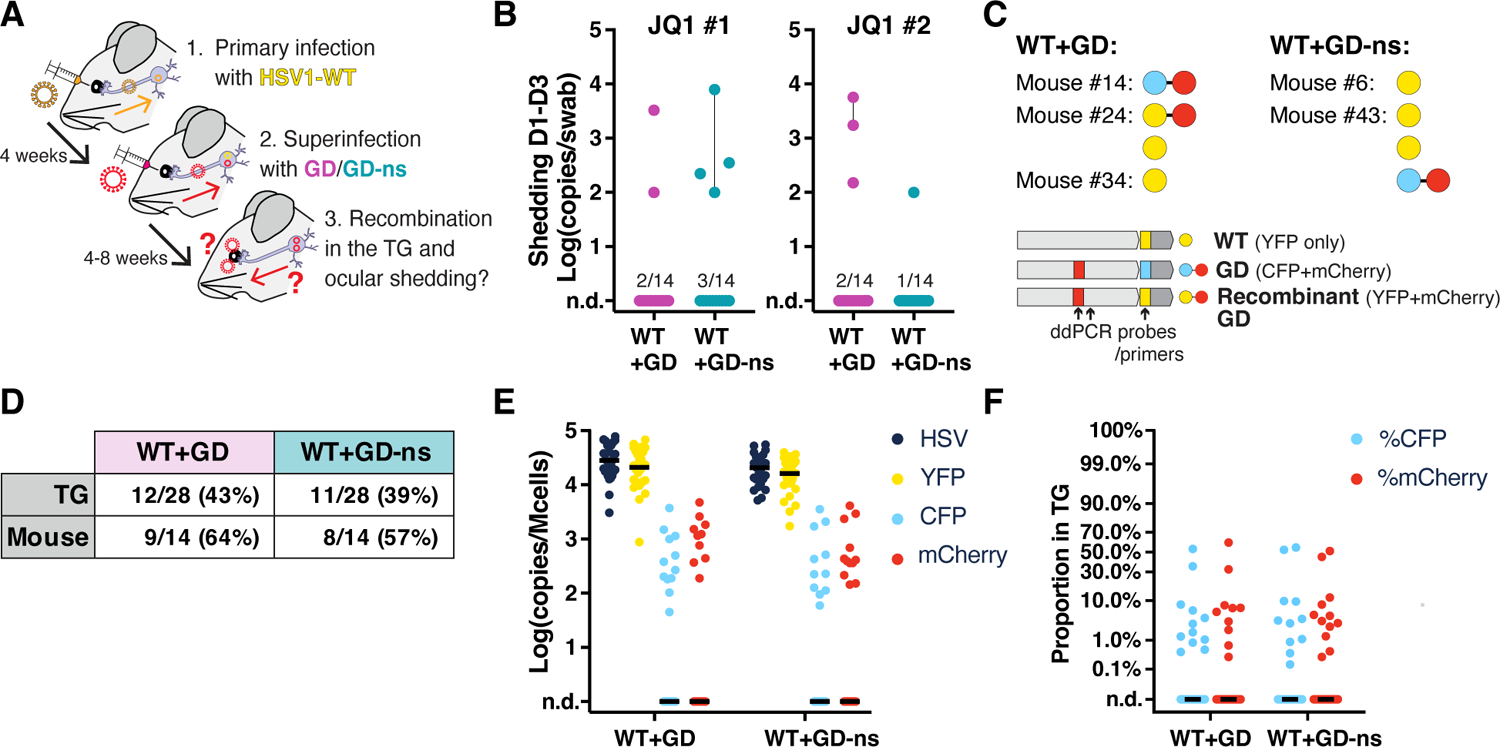
Gene drive spread during latent infection in Swiss-Webter mice. **A**. Experimental outline: Swiss-Webster mice were infected with 10^5^ PFU of HSV1-WT on both eyes after corneal scarification. Four weeks later, mice were superinfected with 10^7^ PFU of GD or GD-ns on both eyes, after corneal scarification. Another four weeks later, latent HSV-1 was reactivated twice with JQ1, two weeks apart. n=14 mice per group. **B**. Titer and number of shedding events in eye swabs on days 1-3 following JQ1 treatment, by qPCR. Shedding events from the same mouse are connected by a line. **C.** Genotyping of positive eye swabs from five mice, using two duplex ddPCR assays. The first assay detected and quantified mCherry levels. The second assay distinguished between YFP and CFP. n=8. **D.** Number and proportion of TG and mice with detectable CFP. **E.** Latent viral load in the TG by duplex ddPCR, detecting mCherry, YFP, CFP markers, or all HSV sequences. n=28. **F.** Proportion of CFP and mCherry in the TG. n=28. Titers are expressed in log-transformed copies per swab, or per million cells after normalization with mouse RPP30 levels. Black lines indicate the median. n.d.: non-detected.

**Figure 8:**
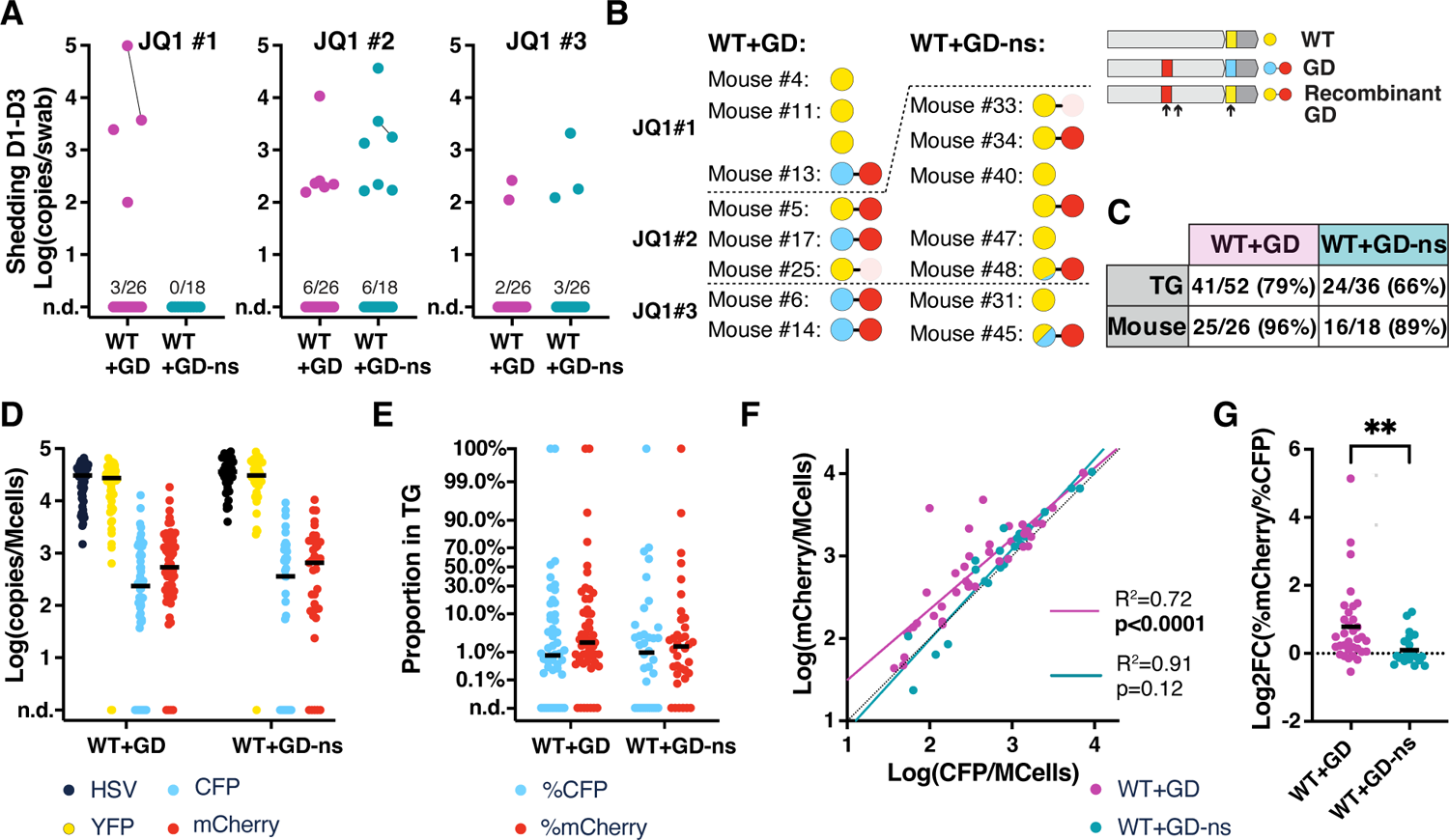
Gene drive spread during latent infection in C57Bl/6 mice C57Bl/6 mice were infected with 10^6^ PFU of HSV1-WT on both eyes after corneal scarification. Four weeks later, mice were superinfected with 10^7^ PFU of GD or GD-ns on both eyes, after corneal scarification. After another four weeks, latent HSV-1 was reactivated three times, two weeks apart, with JQ1 and Buparlisib. n=26 for GD, n=18 for GD-ns. **A**. Titer and number of shedding events in eye swabs on days 1-3 following JQ1 treatment, by qPCR. Shedding events from the same mouse are connected by a line. **B.** Genotyping of positive eye swabs, detecting mCherry, YFP and CFP markers. Light red indicates low mCherry levels, less than 5% of the total swab titer. Swab genotypes were assessed by ddPCR and confirmed by qPCR for low-titer swabs. Details are shown in Supplementary Fig. S12. **C.** Number and proportion of TG and mice with detectable CFP marker from GD/GD-ns. **D-E.** Latent viral load and proportion in the TG. n=52 for GD, n=36 for GD-ns. Black lines indicate the median **F.** mCherry as a function of CFP, and best line fit. GD datapoints were significantly higher than the identity line, suggesting gene drive propagation in the TG. Same data as panel D, excluding non-detected samples. n=38 for GD, n=23 for GD-ns. **G.** Log2 fold-change between the proportion of latent mCherry and CFP, using data from panel E. Samples with low levels of less than 0.5% were excluded. Black lines show the average. Asterisks summarize the results of Welch’s t-test (p=0.0053). n=33 for GD, n=20 for GD-ns. Titers are expressed in log-transformed copies per swab, or per million cells after normalization with mouse RPP30. n.d.: non-detected.

To establish a latent infection, Swiss-Webster mice were infected ocularly with HSV1-WT, with 10^5^ PFU in both eyes after corneal scarification. Four weeks later, animals were inoculated with GD or control GD-ns after corneal scarification (Fig. 7A, Supplementary Fig. S10A, S10B). Immune responses induced by the primary infection limit the spread of a superinfecting virus, allowing mice to be safely inoculated with 10^7^ PFU per eye of GD or GD-ns while being transiently immunosuppressed with the glucocorticoid dexamethasone.

After four more weeks and once the second infection had resolved, latent HSV-1-originating from the primary, secondary infection, or both-was reactivated twice by treating animals with the bromodomain inhibitor JQ1. Mice do not naturally reactivate HSV-1, but JQ1 induction reproducibly induces viral shedding (32, 33). Eye swabs collected on days 1-3 following JQ1 treatment were screened for viral DNA by qPCR. We observed overall low shedding rates. A total of 10 shedding events were detected, with some animals shedding on consecutive days (Fig. 7B). Gene drive-directed recombination was measured in positive swabs using two duplex digital droplet (dd)PCR assays, that distinguish between the YFP, CFP and mCherry markers of WT and gene drive viruses and quantify the proportion of gene drive recombinants. Eight out of ten positive swabs could be successfully genotyped, with the gene drive sequence detected in three of them (Fig. 7C, Supplementary Fig. S10G).

Critically, one of the reactivated viruses (from mouse #24) was a recombinant gene drive virus carrying YFP and mCherry markers, whereas two others carried CFP and mCherry, representing the original gene drive virus. Despite the limited number of shedding events, this showed that GD and GD-ns could successfully reach the latent reservoir and later reactivate, with one recombination event detected.

Next, both TG from each animal were collected and latent viral loads were measured by duplex ddPCR. The CFP marker originating from the superinfecting GD or GD-ns viruses could be detected in around 40% of TG, with 60% of mice having detectable CFP in at least one TG (Fig. 7D). When detected, GD and GD-ns viral loads were approximately two orders of magnitude lower than WT, as measured by the respective titers of YFP and CFP/mCherry (Fig. 7E). Overall, GD and GD-ns represented between 0 and 60% of the total latent viral load (average around 5%, median at 0%), with most detected samples ranging from 1% to 10% (Fig. 7F). This low proportion contrasted with the relatively high frequency of gene drive sequences detected in reactivated swabs (3/8, or 37%), suggesting that the superinfecting GD/GD-ns viruses could successfully reactivate and shed despite representing only a small proportion of the latent reservoir.

Swiss Webster mice are highly susceptible to HSV-1 infection. During primary infection, mice exhibited moderate to severe symptoms, with often extended facial lesions. As a result, most animals had residual scar tissue on their eyes once the primary infection had resolved. Symptoms and final eye scarification were scored during the primary infection, and mice separated into groups with homogenous symptoms and eye scores before superinfection with GD or GD-ns (Supplementary Fig. S10C and S10D). Importantly, at the end of the experiment, GD and GD-ns were detected almost exclusively in the TG of mice with perfect eyes, with even low levels of scarification preventing a successful superinfection (Supplementary Fig. S10E). HSV-1 typically does not cause long-lasting scars in humans, and this residual scar tissue represented an unfortunate confounding factor in our mouse model.

To alleviate this effect, we repeated the experiment in C57Bl/6 mice (Fig. 8). C57Bl/6 are more resistant to HSV-1 infection and typically experience minimal symptoms. C57Bl/6 mice were infected ocularly with HSV1-WT, with 10^6^ PFU in both eyes after corneal scarification.

Compared to Swiss Webster and despite the higher dose, mice exhibited limited symptoms and reduced mortality during primary infection. Around 20% of mice had residual scar tissues, usually on only one eye (Supplementary Fig. S11A-S11D). Four weeks later, mice were superinfected with 10^7^ PFU per eye of GD or GD-ns, while being transiently immunosuppressed with dexamethasone and tacrolimus. Then, another four weeks later, mice were injected with JQ1 and Buparlisib to reactivate latent HSV-1. Buparlisib is a phosphoinositide 3-kinase inhibitor and evidence suggested that it could improve HSV-1 reactivation ((34, 35) and Supplementary Fig. S11E). Reactivation rates ranged from 0 to 33% (Fig. 8A). Most events were close to the detection limit (around 100 copies/swab) and we could successfully obtain viral genotypes from 17 out of 22 reactivation events (Fig. 8B, Supplementary Fig. S12). Nine out of 17 swabs (53%) were either recombinant viruses carrying mCherry and YFP, or original gene drive viruses with mCherry and CFP. In addition, two swabs were genotyped as predominantly WT (from mice #25 and #33), but had detectable amounts of mCherry, representing less than 5% of the total titer (Fig. 8B, Supplementary Fig. S12). In one positive swab with a low viral titer (mouse #45), both YFP and CFP markers, originating from the same *US1/US2* locus, were detected, suggesting a mix of reactivated viruses. Importantly, we detected recombinants expressing YFP and mCherry in mice superinfected with either GD (1/9) or GD-ns (3/8), showing that both viruses could recombine with HSV1-WT. Because of the overall low number of reactivation events, further statistical analysis could not be conducted. In summary, we found that gene drive viruses represented more than 50% of reactivation events, with several examples of recombination with HSV1-WT.

Next, TG were dissected and latent viral loads were measured by duplex ddPCR. GD and GD-ns viruses could be detected in around 79% and 66% of TG, respectively, with more than 90% of mice having detectable GD or GD-ns in at least one TG (Fig. 8C). GD and GD-ns titers were one to two orders of magnitude lower than WT (Fig. 8D), and represented a small percentage of the total latent viral load, ranging from 0.5% to 50% for most samples (average at 10% and median around 1%, Fig. 8E). We investigated whether we could detect evidence of gene drive spread in the TG. In particular, we determined whether mCherry was significantly overrepresented compared to the CFP baseline, as the gene drive cassette containing mCherry potentially recombined with HSV1-WT and increased in frequency. Of note, we observed no correlation between the viral loads in the left and right TG collected from the same animals (Pearson correlation, r^2^=0.11), which allowed us to treat all samples as independent replicates in these analyses (Supplementary Fig. S13A). When plotting the titers of mCherry versus CFP, we observed that GD datapoints were significantly above the identity line (p<0.0001), while GD-ns datapoints were mostly situated along the line and did not significantly deviate from it (Fig. 8F, statistical analysis in Supplementary Fig. S13B). This suggested that the gene drive had recombined with wild-type viruses and increased in frequency in the TG of GD-superinfected mice. To quantify this enrichment, we calculated the fold change between the proportion of mCherry and CFP in the TG (Fig. 8G). On average, in the GD-superinfected samples, we found a 70% increase in the relative proportion of mCherry compared to CFP, which was significantly higher than the control samples with GD-ns where no enrichment was observed (p=0.0053, t-test). In the most extreme cases, the gene drive had spread more than 10-fold over the CFP baseline, for example increasing from 9% to 90% in one sample, or from 1.3% to 10% in another (Supplementary Fig. S13C). Together, this analysis showed a limited but statistically significant spread of the gene drive in the latent wild-type population.

In summary, we found that a superinfecting gene drive virus could reach the latent reservoir and spread in the wild-type population. The drive remained limited in our model and did not reach full penetrance. However, in both Swiss-Webster and C57Bl/6, GD and GD-ns were detected in more than 50% of shedding events after JQ1 reactivation. This indicated that the superinfecting GD and GD-ns viruses could successfully reactivate and shed despite representing only a small proportion of the latent reservoir.

## Discussion

In this manuscript, we designed a gene drive against HSV-1 and showed that it could spread efficiently *in vitro* and *in vivo*. In particular, we observed high levels of co-infection and gene drive-directed recombination in neuronal tissues during herpes encephalitis in mice.

Furthermore, we found evidence that a superinfecting gene drive virus could recombine with HSV1-WT during latent infection. Altogether, this work presented an important proof-of-concept and showed that a gene drive could spread *in vivo* during acute and latent infection.

Our *in vitro* results recapitulated our previous work with hCMV and aligned with the findings of a recent study (11, 28). This showed that a gene drive could be designed in a second herpesvirus with very different infection dynamics. These findings suggest that viral gene drives could be developed in a wide variety of herpesviruses, which significantly expands the potential of the technology.

The spread of a viral gene drive relies on the co-infection of cells by wild-type and engineered viruses. High co-infection levels can easily be achieved in cell culture, either by using cell lines naturally susceptible to co-infection such as N2a cells, or by infecting cells at a high MOI to bypass restriction mechanisms (28). Indirect observations in animal studies and recombination patterns in HSV strains circulating in humans indicate that co-infection events take place *in vivo* (*15–22*). However, whether these co-infection events occur at a high frequency was unknown. Here, we showed that a gene drive could spread efficiently during acute infection in mice, with the level of gene drive-directed recombination increasing from 15% to 80% in some brain regions after only four days. Using fluorescent reporters, we directly observed high levels of co-infection in the TG and the brain, with more than 50% of cells infected by two or more virions in some regions. These findings revealed that HSV-1 achieves high rates of co-infection and recombination during viral spread. Our study highlighted interesting differences depending on the propagation route. For example, co-infection and recombination were low in regions accessed by the optic nerve, and much higher in regions associated with the oculomotor and trigeminal nerves (Fig. 4-6). These observations are intriguing and will be the subject of further studies.

Herpes encephalitis is a severe but rare condition, and HSV disease is more often characterized by chronic lesions on the facial and genital area. We showed that a gene drive virus could reach the latent reservoir and spread in a limited manner in the wild-type population (Fig. 7 and 8). Importantly, gene drive viruses represented around half of the shedding events after drug-induced reactivation, despite these viruses representing a much smaller proportion of the latent viral load. This finding has important implications. It suggests that shedding viruses originated from only a small fraction of the latent reservoir and that a superinfecting gene drive could efficiently reach the physiologically relevant neurons that seed viral reactivation. Mice do not shed HSV-1 spontaneously and our study was limited by the low reactivation rates. Further studies in animals that better recapitulate human disease, such as rabbits or guinea pigs, will be necessary to thoroughly investigate the potential of a gene drive during chronic infection.

This research represents an important milestone toward therapeutic applications. Our future work will focus on designing gene drives that can limit infectivity and reduce disease severity. Interestingly, we noticed that the titer of reactivated gene drive viruses was significantly lower than the titer of wild-type ones (Supplementary Fig. S12C). Insertion of the gene drive sequence did not affect infectivity *in vitro* or during acute infection. However, this unexpected observation suggested that the gene drive may have hampered reactivation or shedding. This could occur either directly if insertion of the gene drive had genetically impaired HSV-1, or indirectly through immune or other host-mediated effects. Similar approaches, where the gene drive virus can reach the latent reservoir efficiently but then suppress reactivation, will form the basis of future strategies. Together, our work may pave the way toward new therapeutics for HSV diseases.

As a final note, the development of gene drives in mosquitoes and other insects has generated important ecological and biosafety concerns (14, 36). Our approach followed the guidelines established by the NIH and the National Academy of Science (37, 38). In the future, the risks and benefits of viral gene drives will need to be properly addressed and discussed with the scientific community.

## Materials and Methods

### Cells and viruses

African green monkey epithelial Vero cells and murine neuroblastoma N2a cells were obtained from the ATCC and cultured in DMEM (Corning, Corning, NY, USA) supplemented with 10% FBS (Sigma-Aldrich, St-Louis, MO, USA). Cells were maintained at 37 °C in a 5% CO_2_ humidified incubator and frequently tested negative for mycoplasma contamination.

Unmodified HSV-1 strain 17+ and HSV1-CFP expressing cyan fluorescent protein mTurquoise2 from the US1/2 locus (15) were provided by Matthew Taylor (Montana State University, USA). Viruses generated for this study were made by modifying HSV-1 and HSV1-CFP, as described below. To prepare viral stocks for cell culture experiments, Vero cells in 15 cm dishes were infected for one hour at MOI=0.01, and kept in culture for 48 hours or until 100% cytopathic effect was observed. Cells and supernatant were scraped out of the plate, sonicated three times at maximum power with a probe sonicator, and debris pelleted away by centrifugation (2000 rpm, 10 minutes, 4 °C). Media containing viruses was collected in single-use aliquots and titers measured by plaque assay.

For high-titer and high-purity viral stocks used for animal experiments, Vero cells in 15 cm dishes were infected for one hour at MOI=0.01, and kept in culture for 48 hours or until 100% cytopathic effect was observed. Supernatants and cells were collected, and cells were pelleted by centrifugation (2000 rpm, 5 minutes, 4 °C). Supernatants were collected in clean tubes and reserved for later. Cell pellets were resuspended in a small volume of culture media and cell-bound virions were released by two cycles of freeze-thaw in dry ice. Debris were pelleted again and the supernatant containing released virions was combined with the supernatant reserved earlier. Virions were then pelleted by ultracentrifugation (22,000 rpm, 90 min, 4 °C, Beckman-Coulter rotor SW28) on a 5-mL cushion of 30% sucrose. Supernatants were discarded, and virions were resuspended in PBS containing 2% BSA. Single-use aliquots were prepared and titers were measured by plaque assay.

Co-infection experiments were performed in 12-well plates by co-infecting N2a cells with HSV1-WT and gene drive viruses for 1 h, with a total MOI of 1, before replacing inoculum with 1mL of fresh medium. 100uL of supernatant was collected at regular intervals and analyzed by plaque assay.

### Cloning and generation of recombinant viruses

A donor plasmid containing the gene drive cassette against the HSV-1 *UL37-38* intergenic region (GD and derivatives) was generated by serial modifications of the GD-mCherry donor plasmid used in our previous study (11). All modifications were carried out by Gibson assembly (NEB, Ipswich, MA, USA), using PCR products from other plasmids or synthesized DNA fragments (GeneArtTM StringTM fragments, ThermoFisher, USA). The final GD donor plasmid included homology arms for the *UL37-38* region, the CBH promoter driving *SpCas9* followed by the SV40 polyA terminator, the CMV promoter driving an *mCherry* reporter followed by the beta-globin polyA signal, and the U6 promoter controlling gRNA expression.

The functional GD plasmid carried a gRNA targeting the *UL37-38* region (ACGGGATGCCGGGACTTAAG), while the non-specific GD-ns control targeted a sequence absent in HSV-1 (ACATCGCGGTCGCGCGTCGG). GD-i1Cas9 donor construct was subsequently generated by removing *SpCas9* by digestion and ligation. Donor constructs to insert CMV-driven yellow (YFP) or red (RFP) fluorescent protein reporters into the *US1/US2* locus were built similarly, by replacing *mTurquoise* with *mCitrine2* or *mScarlet2* in a donor plasmid for the US1/US2 region, respectively (pGL002, from ref (15)). Of note, the YFP, CFP and RFP reporters carried a nuclear localization signal.

To build recombinant viruses, 1.5 million Vero cells were co-transfected with linearized donor plasmids and purified HSV-1 strain 17+ or HSV1-CFP viral DNA. Viral DNA was purified from infected cells by HIRT DNA extraction, as described previously (11). Transfection was performed by Nucleofection (Lonza, Basel, Switzerland) and cells were plated in a single 6-well. After 2-4 days, mCherry-expressing viral plaques were isolated and purified by several rounds of serial dilutions and plaque purification. Purity and absence of unmodified viruses were assayed by PCR and Sanger sequencing after DNA extraction (DNeasy kit, Qiagen, Germantown, MD, USA). Viral stocks were produced as specified above and titered by plaque assay.

### Plaque assay

Plaque assays were performed either directly from cell culture supernatants, or from frozen mouse tissues. To release infectious virions from tissues, frozen samples were resuspended in cell culture media and disrupted using a gentle tissue homogenizer (Pellet Pestle, Fisher Scientific, USA). Samples were sonicated three times at maximum power with a probe sonicator, and debris pelleted away by centrifugation (2000 rpm, 10 minutes, 4 °C). Volumes were adjusted to a final volume of 1 mL, and titers were measured by plaque assay.

Viral titers and recombination levels were determined by plaque assay with 10-fold serial dilutions. Confluent Vero cells in 24-well plates were incubated for 1 h with 100uL of inoculum, and overlaid with 1mL of complete media containing 1% methylcellulose, prepared using DMEM powder (Thermo) and Methylcellulose (sigma). After two or three days, fluorescent plaques expressing YFP, CFP and/or mCherry were manually counted using a Nikon Eclipse Ti2 inverted microscope. Every viral plaque was analyzed on both YFP, CFP and red channel. 5-100 plaques were counted per well, and each data point was the average of 3-4 technical replicates (i.e., 3-4 different wells). Images of fluorescent viral plaques were acquired with an EVOS automated microscope and adjusted for contrast and exposure with ImageJ (v2.1.0). The deconvolution of Sanger sequencing in Fig. 3F was performed using Synthego ICE online tools (https://ice.synthego.com).

### Mouse experiments

All animal procedures were approved by the Institutional Animal Care and Use Committee of the Buck Institute and Fred Hutchinson Cancer Center. This study was carried out in strict accordance with the recommendations in the Guide for the Care and Use of Laboratory Animals of the National Institutes of Health (“The Guide”).

### Acute infection after intravitreal inoculation

Acute infections were performed at the Buck Institute. Male and female Balb/c between five and eight weeks-old were infected by intravitreal injection in the left eye, as described previously (39, 40). Briefly, mice were anesthetized by intraperitoneal injection of ketamine (100 mg/kg) and xylazine (10 mg/kg) and laid prone under a stereo microscope. The left eye was treated with a thin layer of veterinary ophthalmic ointment and the sclera was exposed using an ophthalmic forceps. 2 µl of HSV-1 stock containing 10^6^ pfu was injected slowly in the intravitreal space, using a 5uL Hamilton syringe and a 30 gauge needle. In the days following infection, mice were treated with sustained-release buprenorphine to minimize pain (Ethiqa XR, Fidelis Animal Health, North Brunswick, NJ, USA). Animals were humanely euthanized after two to four days. For plaque assay analysis, tissues were collected and snap-freezed in liquid nitrogen.

### Latent infection after corneal scarification

Latent infections were performed at the Fred Hutch Cancer Center, using female Swiss-Webster or C57bl/6 mice five to six weeks-old purchased from Taconic (Germantown, NY, USA). Mice were anesthetized by intraperitoneal injection of ketamine (100 mg/kg) and xylazine (10 mg/kg) and laid under a stereo microscope. Mice corneas were lightly scarified using a 28-gauge needle, and 4uL of viral inoculum dispensed on both eyes. Swiss-Webster and C57bl/6 mice were infected with 10^5^ and 10^6^ PFU, respectively. Following inoculation, ophthalmic drops of local analgesic (Diclofenac) were deposited on both eyes, and the analgesic Meloxicam was added to the drinking water *ad libitum* for 1-5 days following infection. From five to fifteen days following primary infection, symptoms of infection were reported and scored using an in-house scoring system. Mice experiencing severe symptoms were humanely euthanized. Once the infection had fully resolved, final eye scarification levels were scored in both eyes and averaged, using the following scores: 0: perfect eye; 1: lightly damaged and/or cloudy cornea, 2: scar tissue covering a small portion of the eye; 3: scar tissue covering most of the eye; 4: extremely bad looking eye, fully blind.

The second infection with GD and GD-ns was performed the same way four weeks after the primary infection, with 10^7^ PFU per eye. Mice were transiently immunosuppressed with dexamethasone (Fig. 7 in Swiss-Webster) or with dexamethasone and tacrolimus (Fig. 8 in C57bl/6). Tacrolimus and Dexamethasone were diluted in the drinking water and administrated *ad libitum* from one day before to seven days after infection. Drug concentration was calculated according to the average mice weight, considering that mice drink around 5mL per day, in order to reach a dose of 1 mg/kg/day and 2 mg/kg/day for dexamethasone and tacrolimus, respectively. For example, for an average mouse weight of 25g, dexamethasone and tacrolimus were diluted at 5mg/L and 10mg/L, respectively. Of note, drugs were diluted in medidrop sucralose (ClearH20, Westbrook, ME, USA) instead of regular drinking water, to make tacrolimus more palatable to mice.

### HSV reactivation

HSV reactivation was performed by intraperitoneal injection of JQ1 (MedChemExpress, NJ, USA) at a dose of 50 mg/kg, and, when indicated, Buparlisib (MedChemExpress, NJ, USA) at a dose of 20 mg/kg. JQ1 and Buparlisib were prepared from stock solutions (10x at 50mg/mL and 100x at 200mg/mL, in DMSO, respectively) by dilution in a vehicle solution of 10% w/v 2-hydroxypropyl-!3-cyclodextrin (Sigma-Aldrich, St-Louis, MO, USA) in PBS. In the C57bl/6 experiments (Fig. 8), mice were transiently immunosuppressed with dexamethasone and tacrolimus in the drinking water, at the concentrations indicated above, from one day before to three days after JQ1/Buparlisib injection. Mouse eyes were gently swabbed with cotton swabs moistened with PBS, on day one to three following injection. Swabs were collected into vials containing 1 ml of digestion buffer (KCL, Tris HCl pH8.0, EDTA, Igepal CA-630) and stored at 4°C before DNA extraction.

### HSV quantification of viral loads in swabs and tissues

#### HSV quantification of eye swabs by qPCR

DNA was extracted from 200 µl of swab digestion buffer using QiaAmp 96 DNA Blood Kits (Qiagen, Germantown, MD, USA) and eluted into 100 µl AE buffer (Qiagen, Germantown, MD, USA). 10 µl of eluted DNA was used to setup 30 µl real-time Taqman quantitative PCR reactions, using QuantiTect multiplex PCR mix (Qiagen, Germantown, MD, USA), using the following PCR cycling conditions: 1 cycle at 50°C for 2 minutes, 1 cycle at 95°C for 15 minutes, and 45 cycles of 94°C for 1 minute and 60°C for 1 minute. Exo internal control was spiked into each PCR reaction to monitor inhibition. A negative result was accepted only if the internal control was positive with a cycle threshold (CT) within 3 cycles of the Exo CT of no template controls. Primers and probes have been described previously and are provided in Supplementary Table S1 (41). Swabs positive for HSV were then further analyzed by ddPCR, as described below.

#### Duplex digital droplet PCR (ddPCR) of tissues and swabs

Total genomic DNA was isolated from ganglionic tissues using the DNeasy Blood and tissues kit (Qiagen, Germantown, MD, USA) and eluted in 60 µl of EB buffer, per the manufacturer’s protocol.

Quantification of the YFP, CFP and mCherry markers, as well as total HSV viral load was measured with two separate duplex ddPCR, using 10uL of eluted DNA. ddPCR was performed using the QX200 Droplet Digital PCR System and ddPCR Supermix for Probes (No dUTP) from Biorad (Hercules, CA, USA), following the manufacturer’s instructions. Primers were used at a final concentration of 900nM and probes at 250nM (Supplementary Table S1). Primers and probes were ordered from IDT (Coralville, IA, USA), using their custom PrimeTime ZEN double-quenched qPCR probes, with FAM and HEX fluorescent dyes. The first duplex assay used two sets of primers/probes to quantify mCherry (HEX probe) and HSV *UL38* gene (FAM probe). *UL38* primers/probe set was located in the *UL38* viral gene and recognized both wild-type and gene drive genomes. The second duplex assay distinguished between YFP and CFP, using one set of common primers amplifying both markers and YFP and CFP-specific probes with FAM and HEX dyes, respectively. Primer specificity and sensitivity were validated on plasmid DNA before use in mouse samples. A limit of detection of three copies per reaction was used throughout the study, except in Fig. 7D-7F, where a cutoff of 10 was applied to mCherry to account for a small PCR contamination. Final titers were normalized and expressed in log-transformed copies per million cells (MCells). Cell numbers in tissue samples were quantified by ddPCR using a mouse-specific RPP30 primer/probe set (42).

The duplex assays allowed us to determine the proportion of latent mCherry and CFP, using absolute values measured in the same PCR reaction, thus, limiting technical variation.

Proportions were calculated as follows:

*%mCherry = 100 * mCherry/UL38*

*%CFP = 100 * CFP/(CFP + YFP)*

#### Swab genotyping

Positive swabs identified by qPCR were analyzed using the duplex ddPCR assays described above. Samples expressing YFP only with no detectable CFP and mCherry were categorized as wild-type. Swabs expressing mCherry at the same level as *UL38* were categorized as gene drive. Swabs expressing both CFP and mCherry represented the original GD/GD-ns, while swabs expressing both YFP and mCherry represented recombinants (Supplementary Fig. S10G, S12). Some swabs expressed mCherry one to two orders of magnitude lower than HSV and were genotyped as wild-type but with detectable amounts of mCherry. For low-titer swabs, the genotype was further confirmed by duplex qPCR using the same primers.

### Brain and TG imaging and image analysis

#### Tissue processing and image collection

Balb/c mice were infected ocularly with equivalent amounts of three viruses expressing either YFP, CFP or RFP from the *US1-US2* locus. A total of 10^6^ PFU was inoculated intravitreally in the left eye. Of note. the fluorescent proteins carry nuclear localization signals. They are expressed in infected cells and are not incorporated into virions. Four days after infection, mice were injected intraperitoneally with a terminal dose of euthanasia solution containing Sodium Pentobarbitol (Euthasol). Once unresponsive, mice were subjected to thoracotomy and transcardially perfused with PBS followed by 4% Paraformaldehyde-Lysine-Periodate solution (PLP) through the aorta to fix tissues (43).

Brains, TG, and eyes were dissected, fixed in PLP overnight, and transferred to a 20% sucrose solution for 24 hours, and finally to a 30% sucrose solution for at least 24 hours for cryo-protection. All tissues were stored in 30% sucrose before processing. This protocol was a courtesy of J.P. Card (40).

TG were embedded in OCT and serial sections of 15 µm made using a Cryostat (Zeiss) at −20°C. TG sections were mounted on subbing solution-treated slides and polymerizing mounting media containing DAPI (Vectashield, Vector Labs, Burlingame, CA, USA) was added before coverslipping. Brains were immobilized in 30% sucrose on a freezing microtome stage set to −18° C (Physitemp, Clifton, NJ). Serial coronal sections at 30 µm on a horizontal sliding microtome (AO Optical) were collected. Brain sections were binned into six parallel groups. One bin was arranged and mounted on slides before counterstaining with polymerizing mounting media containing DAPI and coverslipping. Epifluorescence imaging was performed on a Nikon Ti-Eclipse (Nikon Instruments, Melville, NY, USA) inverted microscope equipped with a SpectraX LED (Lumencor, Beaverton, OR, USA) excitation module and fast-switching emission filter wheels (Prior Scientific, Rockland, MA, USA).

Fluorescence imaging used paired excitation/emission filters and dichroic mirrors for DAPI, CFP, YFP and TRITC (Chroma Technology Corp., Bellow Falls, VT, USA). All images were acquired with an iXon 896 EM-CCD (Andor Technology LTD, Belfast, NI, USA) camera using NIS-Elements software. Image tiles with 4x and 10x Phase objectives were acquired to assess fluorescent protein expression across sectioned tissues. Specific regions were imaged with the 20x ELWD to acquire detailed localization and fluorescent protein expression images for subsequent data analysis.

#### Image analysis

From the five animals originally infected, one animal was not included as very few infected cells could be detected in the brain and TG, suggesting that the infection had failed. Furthermore, one brain from the remaining four mice was irremediably damaged during processing. Thus, the analysis was conducted on four TG and three brains.

Using ImageJ (v2.14.0/1.54f), Nd2 images acquired with Nikon NIS-Element software were batch converted into tiff files using an ImageJ macro (modified from https://github.com/singingstars/). During the conversion process, brightfield and DAPI channels were discarded, and ImageJ background subtraction was performed on the YFP, CFP and RFP channels. An additional grayscale channel was created, composed of the maximum projection of the YFP, CFP and RFP channels. This composite channel contained every cell irrespective of the original color and was used for segmentation (Supplementary Fig. S3A). Machine learning-assisted segmentation was performed using an online analysis tool from www.biodock.ai (Biodock, AI Software Platform. Biodock 2023). The software was trained to recognize cells on the gray channel using a few training images, and segmentation was then run on the entire dataset. Around 3-4% of cells with aberrant area or eccentricity were discarded, and average signal intensity was measured in the original YFP, CFP and RFP channels for each detected cell. Of note, for the TG, this analysis was performed using only the YFP and CFP channels. Data was then further processed and plotted using R (RStudio v2023.09.1+494). For YFP and CFP, the intensity was simply log10 converted. Because RFP had a higher background and different intensity ranges across images, RFP intensity was first scaled across images and then log10 converted. Stringent intensity thresholds were applied on the three channels and used to quantify cells infected with one, two, or three viruses, with around 5% of cells below thresholds being discarded (Supplementary Fig. S5B). Thresholds were chosen stringently to unequivocally identify co-infected cells, and are reported in Supplementary Fig. S3C and S5C. For visualization and plotting, signal intensities in the YFP, CFP and RFP channels were converted into CYMK color space. Data is provided in Supplementary files S1 and S2.

Representative images shown in the manuscript were minimally adjusted for contrast and exposure using ImageJ. Some images were rotated or flipped horizontally to consistently present the brain in a caudal direction, with the left hemisphere on the right side.

#### Statistics and reproducibility

Experiments were carried out in multiple replicates. Investigators were blinded when performing plaque assays, collecting swabs, and analyzing DNA samples. No data was excluded, except when indicated in the main text, methods or figure legends. Statistical analyses were performed using GraphPad Prism version 10.1.1 for macOS (GraphPad Software, USA, www.graphpad.com). Statistical tests and their results are described in the text and figure legends.

## Code and data availability

The data supporting the findings of this study are available within the paper and its Supplementary files. Plasmids, viruses, and other reagents developed in this study are available upon request and subject to standard material transfer agreements with the Buck Institute and Fred Hutch Cancer Center. Any other relevant data are available upon reasonable request.

## Acknowledgments

We thank ophthalmologist Koji Kitazawa (Buck Institute and Kyoto Prefectural University of Medicine) for training MW with intravitreal injections. We thank members of the Verdin and Jerome labs for technical and conceptual help. This study was funded through institutional support from the Buck Institute for Research on Aging and the Fred Hutch Cancer Center. In particular, MW received funding from the VIDD faculty initiative award from the Fred Hutch Cancer Center.

## Author contributions

MW designed the study with input from EV, KRJ and MPT. MW performed all *in vitro* experiments. MW and RR wrote the IACUC protocol at the Buck Institute. MW, RR, AKH, PAM, MAL, DES and MA contributed to mouse husbandry and mouse experiments. LMK, LS and TKS processed samples at the University of Washington Virology Laboratory. MW and MPT performed the analysis of co-infection in the brain. MW, MPT, KRJ and EV analyzed the data. MW wrote the manuscript with input from all authors. EV and KRJ supervised and funded the project.

---

**Supplementary Table S1.**
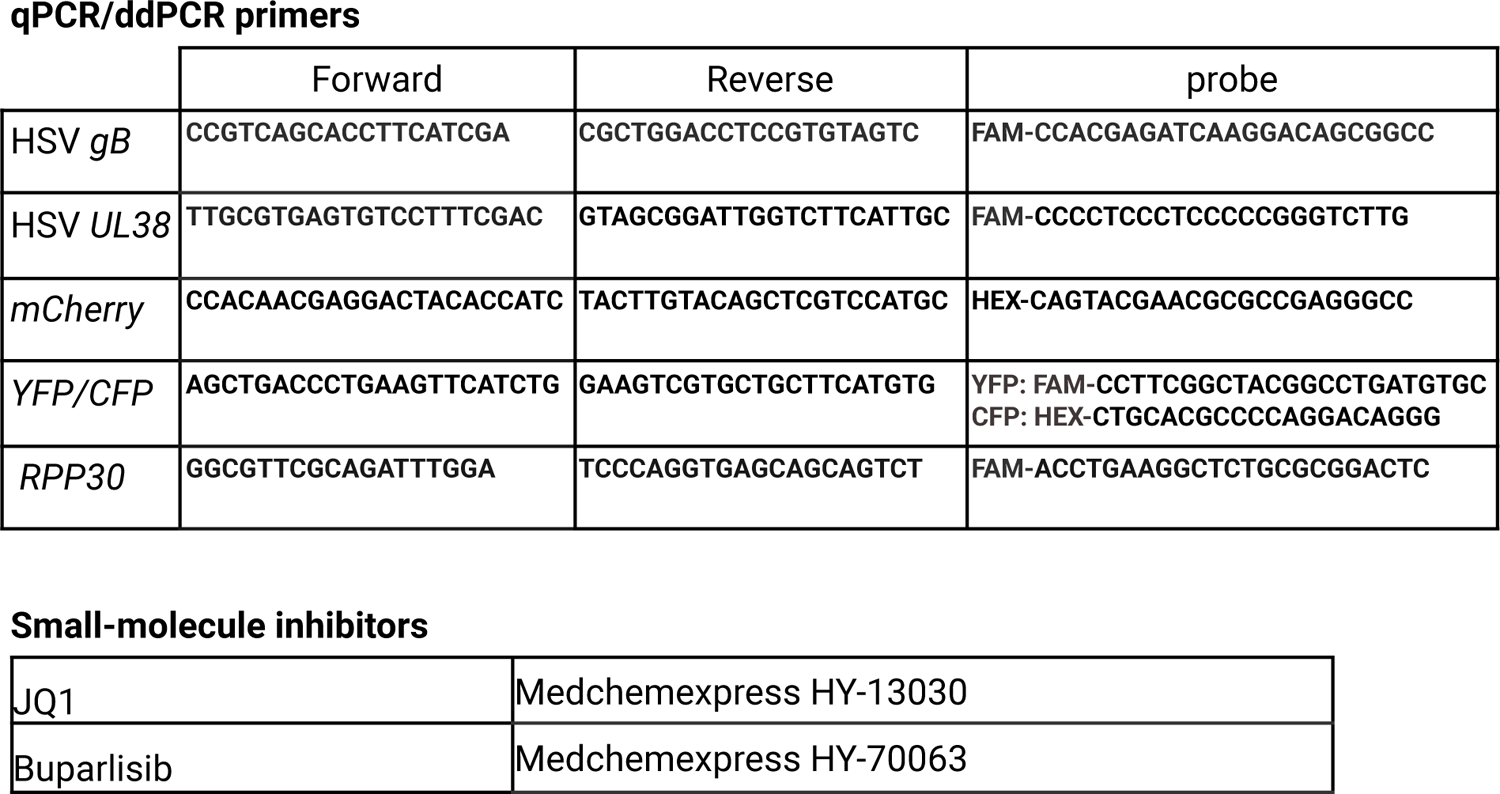

**Supplementary Figure S1.**
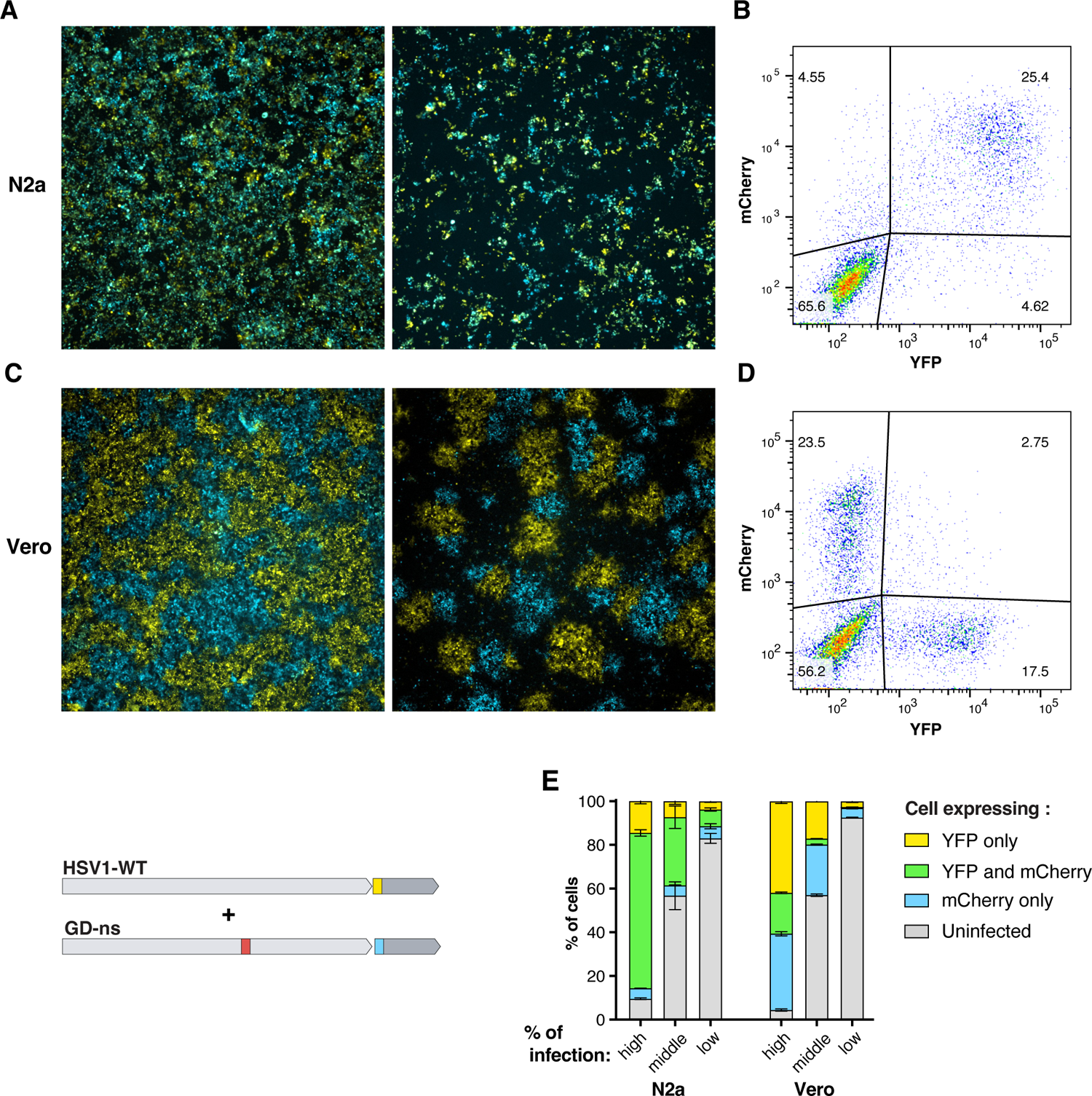
N2a cells sustain high levels of co-infection. N2a and Vero cells were coinfected with WT virus expressing YFP and GD-ns virus expressing CFP and mCherry, at low MOI (from 0.1 to 0.001). Two days later, the proportion of infected and co-infected cells was analyzed by microscopy (A, C) or flow cytometry (panels B, D and E). At both high, middle, or low total infection rates, N2a cells showed a high level of co-infection, while Vero cells had very little co-infection. In panel E, data show mean and SEM between 6 (high co-infection rate) or 3 (middle and low infection rates) biological replicates. High, middle, and low infection rates indicate that around 90%, 40%, or 10% of all cells are infected, respectively.

**Supplementary Figure S2.**
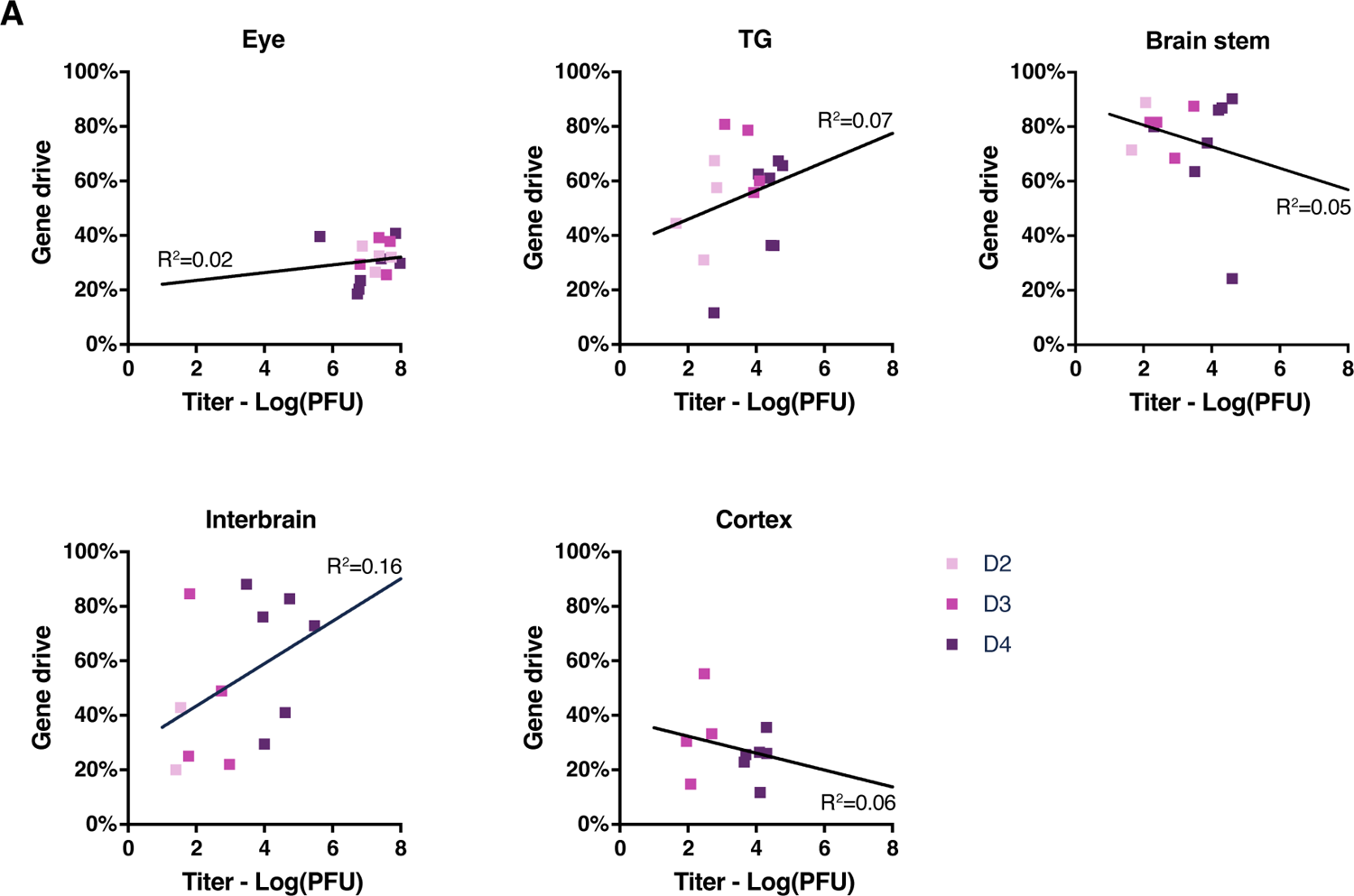
No correlation between viral titers and gene drive-directed recombination. Simple linear regression between viral titers and the level of gene drive-directed recombinants in the different brain regions.

**Supplementary Figure S3.**
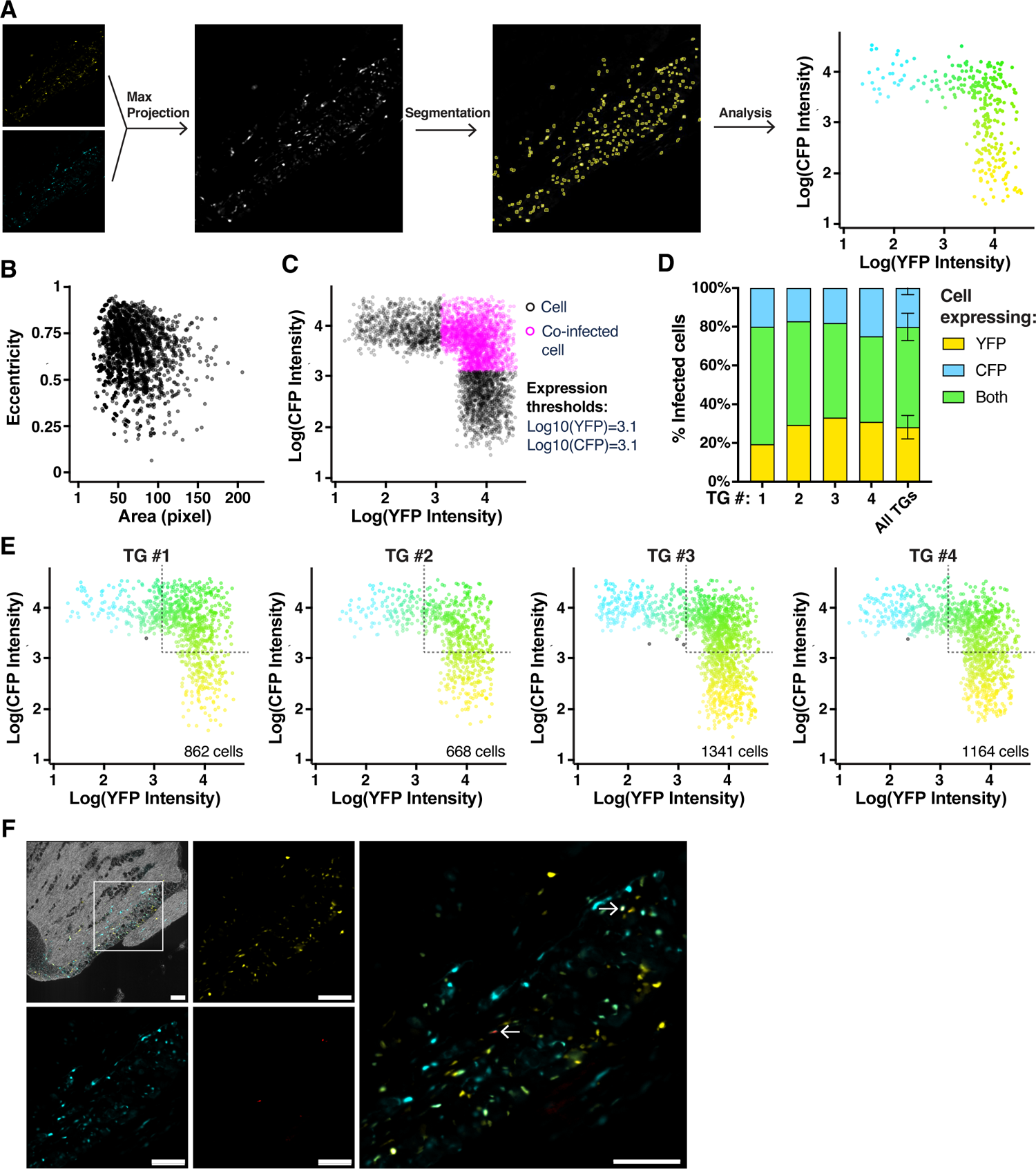
Machine learning-assisted analysis of co-infection in the TG. **A**. Summary of the analysis pipeline: individual color channels were merged to create composite grayscale images. Infected cells were automatically segmented using machine learning analysis software from biodock.ai, allowing quantification of mono- and co-infected cells. 4035 cells were detected, originating from 53 images and n=4 animals. **B**. Cell area and eccentricity after cell segmentation. **C**. Co-infected cells and expression thresholds. **D**. Percentage of infected cells expressing YFP, CFP, or both, in the four biological replicates. **E**. High consistency between biological replicates. **F**. Representative images of TG sections. Scale bars: 100 µm. Arrows indicate cells co-expressing YFP, CFP and RFP together.

**Supplementary Figure S4.**
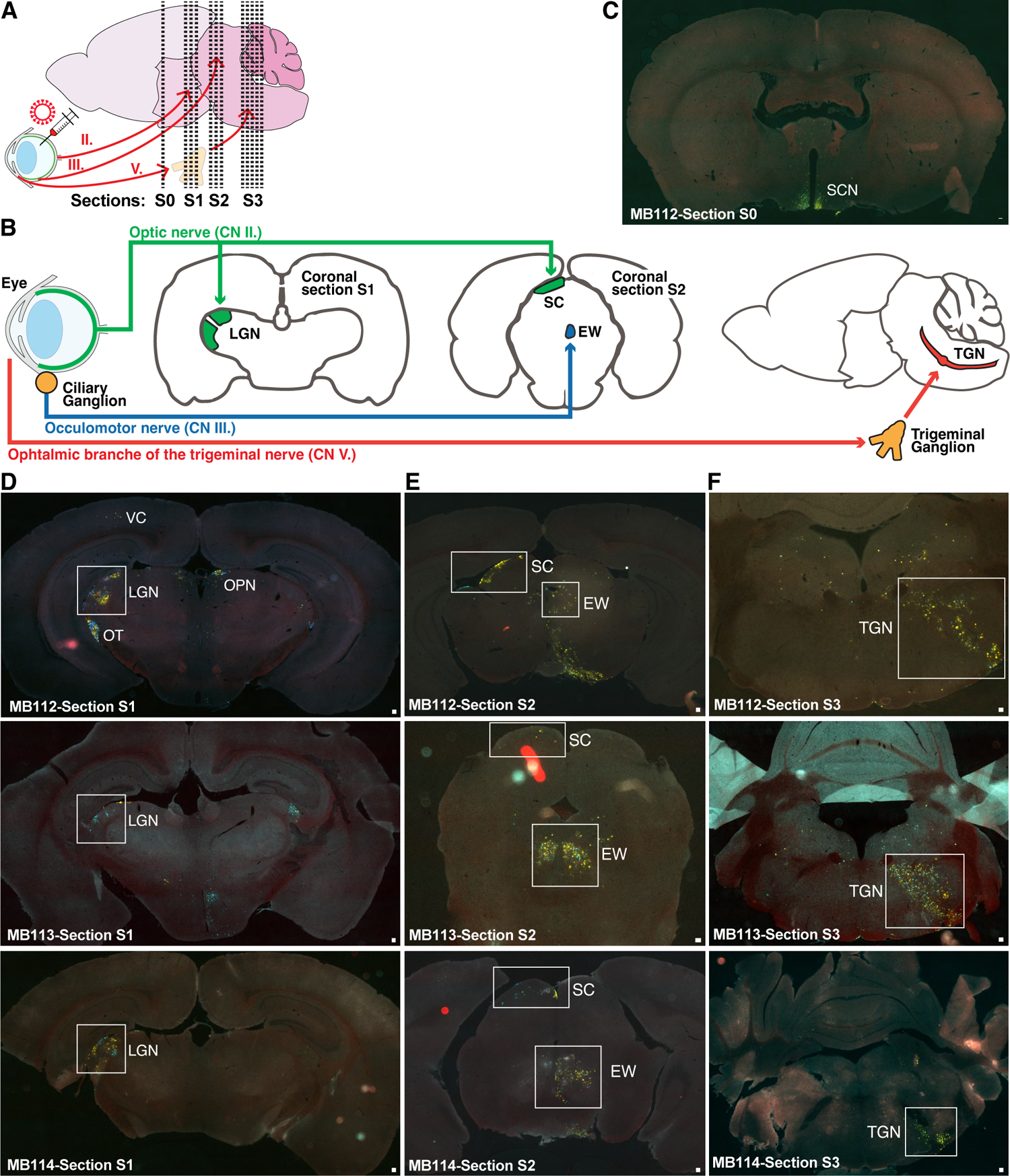
Infection of the visual system in the brain. **A**. Coronal sections of the brain collected after ocular infection. **B**. Regions of primary spread in the visual system. HSV-1 infects retinal neurons and travels via the optic nerve to the LGN in the thalamus and to the SC in the midbrain. After infection of the ciliary ganglion, HSV-1 travels to the EW in the midbrain via the oculomotor nerve. Finally, HSV-1 travels via the trigeminal nerve through the TG, reaching the TGN in the brain stem. **C-F**. Reproducible infection of the visual system, as shown in three biological replicates (brain ID# MB112, MB113 and MB114). SCN: suprachiasmatic nuclei, LGN: lateral geniculate nucleus. OT: optic tract, OPT: olivary pretectal nucleus, VC: visual cortex SC: superior colliculus, EW: Edinger-Westphal nucleus, TGN: trigeminal nerve nuclei. Scale bars: 100 µm.

**Supplementary Figure S5.**
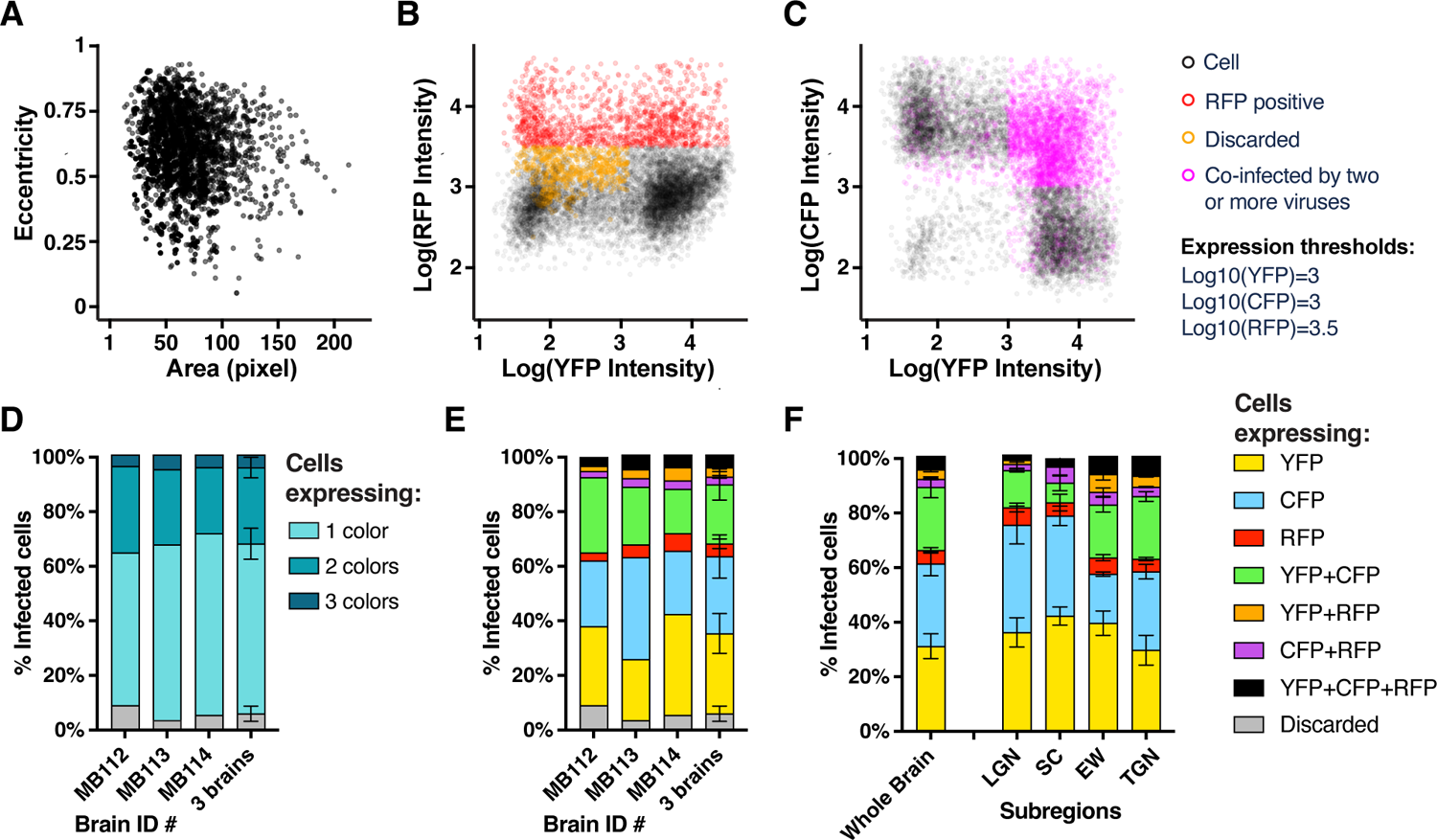
Machine learning-assisted analysis of co-infection in the brain. 10,028 cells were detected, originating from 95 images and n=3 animals (brain ID# MB112, MB113 and MB114). **A**. Cell area and eccentricity. **B**. RFP intensity threshold and discarded cells. **C.** Co-infected cells and expression thresholds **D**. Percentage of infected cells expressing one, two, or three fluorescent markers in the three biological replicates. **E-F.** Percentage of infected cells expressing the different color combinations, in the whole brain and the three biological replicates (E), or across the subregions (F). n=3.

**Supplementary Figure S6.**
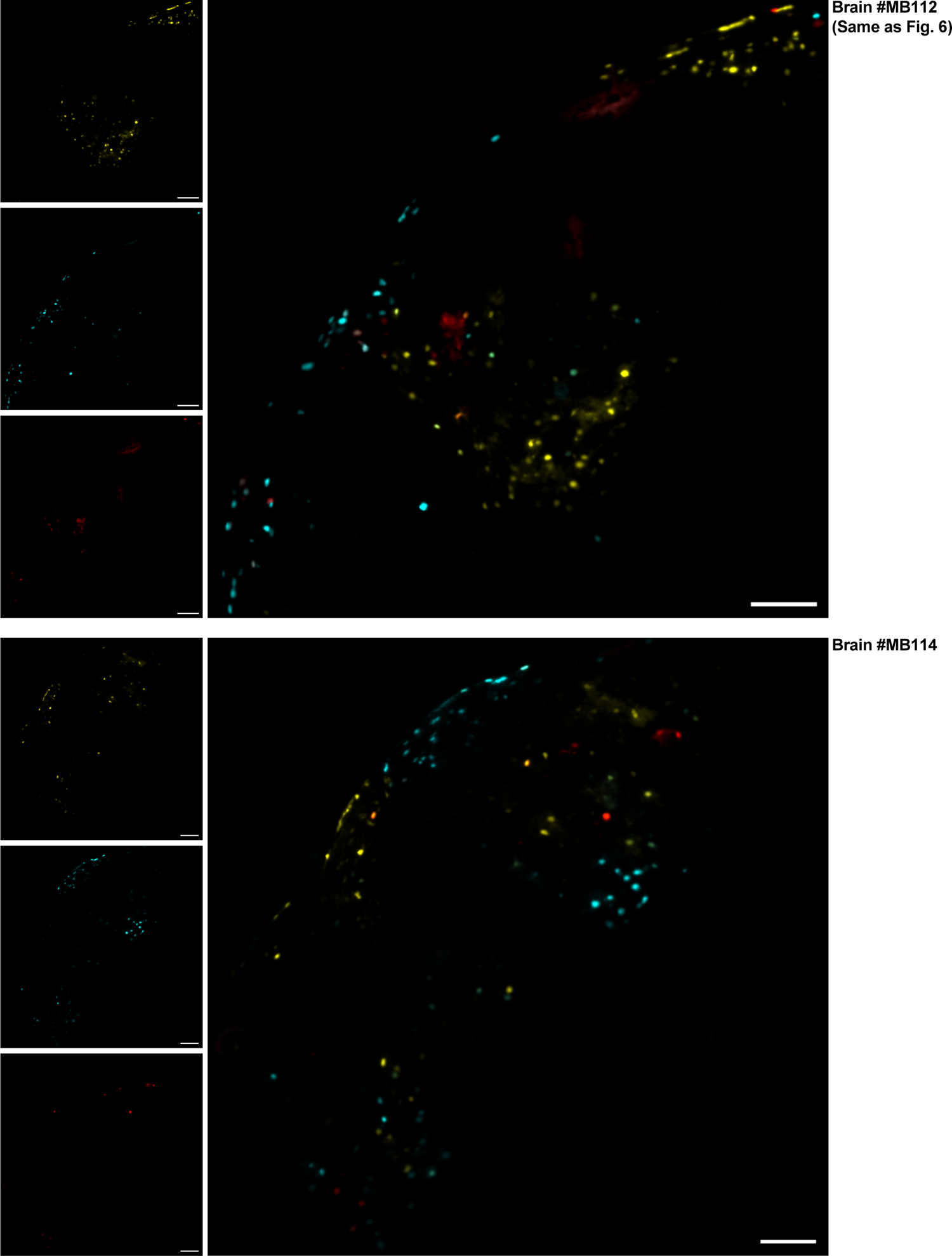
Low co-infection levels in the lateral geniculate nucleus. Representative images of the lateral geniculate nucleus (LGN) from two different biological replicates. Infected cells form tight foci expressing only one color, with co-infected cells at the boundaries. Scale bars: 100 µm.

**Supplementary Figure S7.**
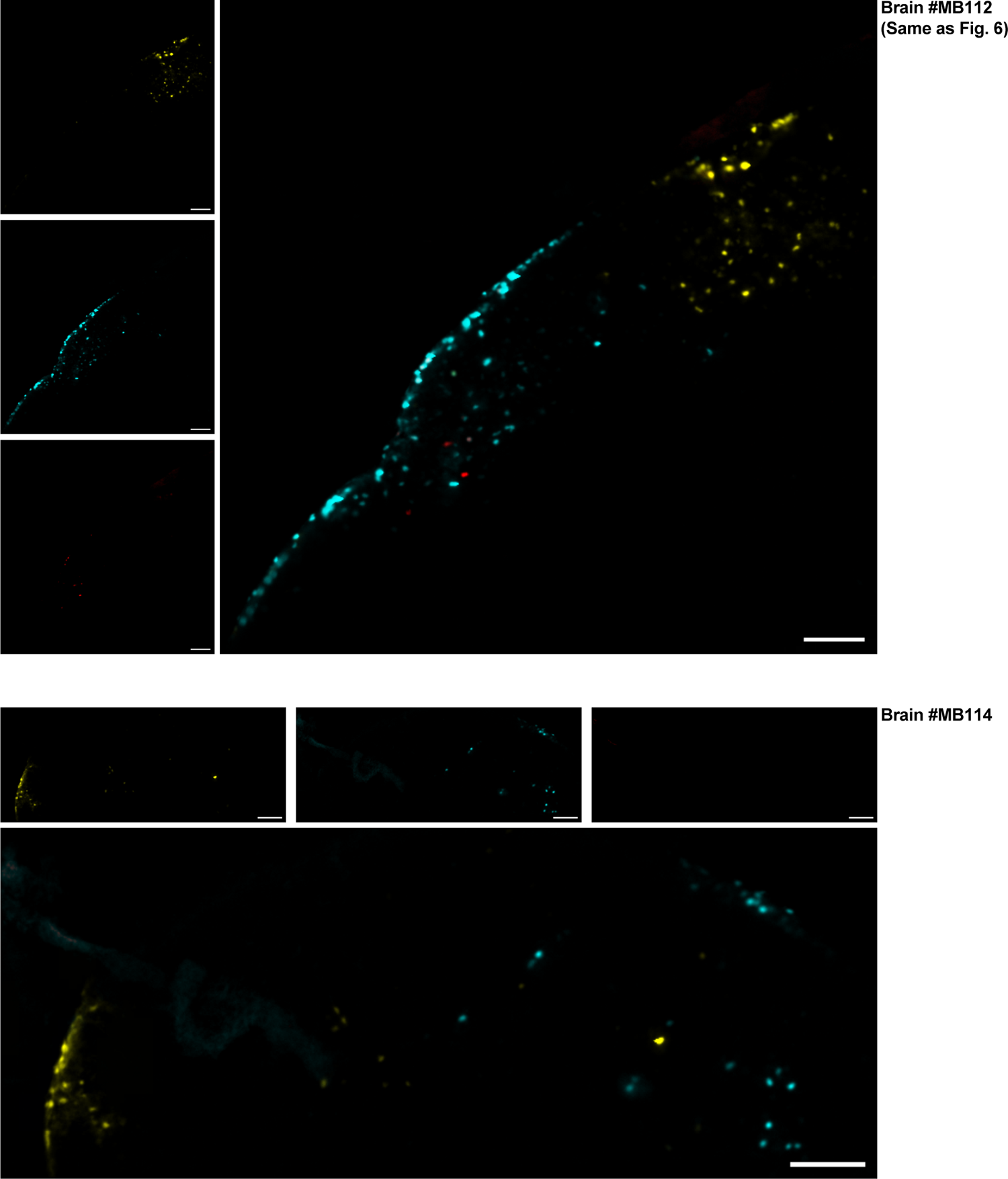
Low co-infection levels in the superior colliculus. Representative images of the superior colliculus (SC) from two different biological replicates. Infected cells form tight foci expressing only one color, with co-infected cells at the boundaries. Scale bars: 100 µm.

**Supplementary Figure S8.**
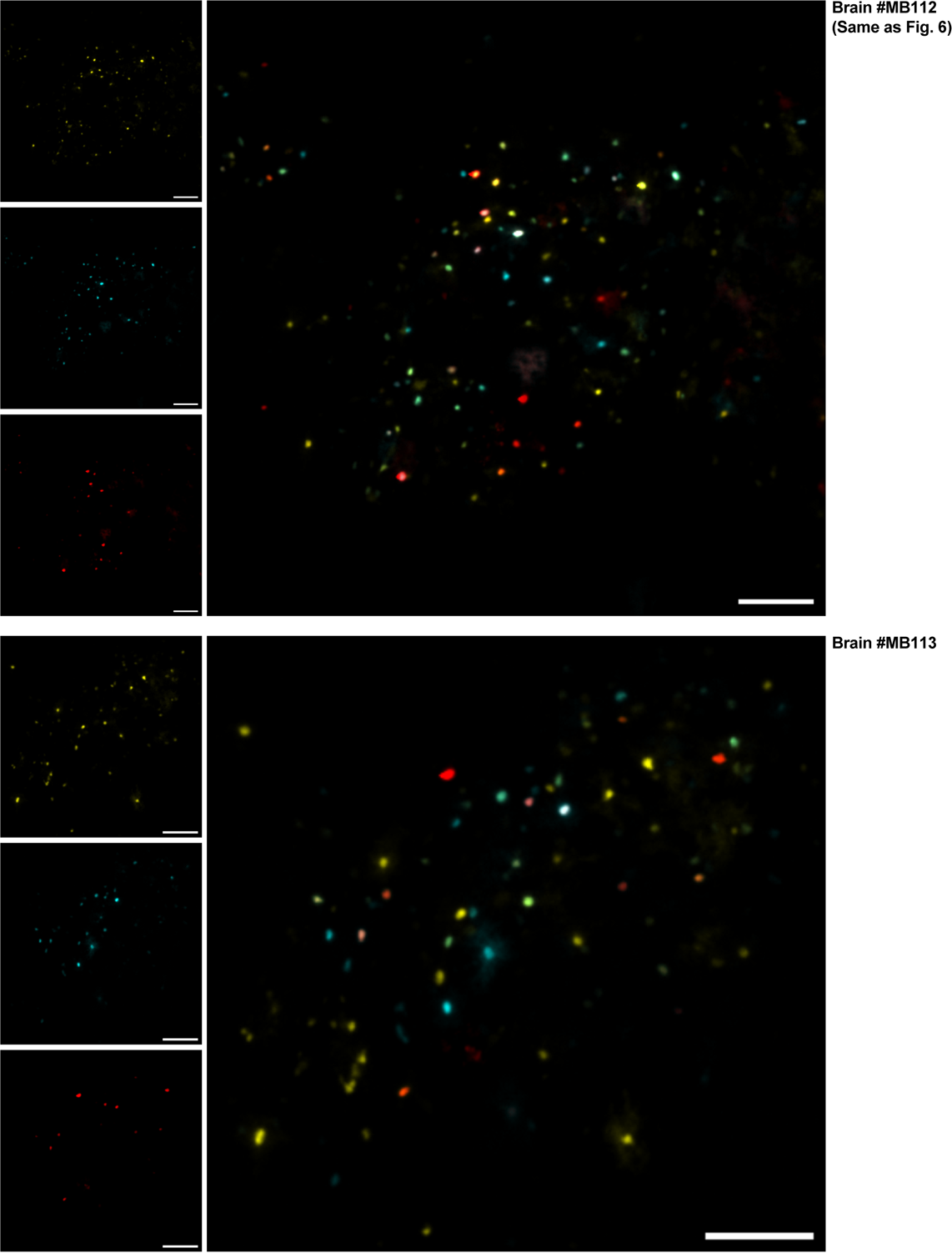
High co-infection levels in the Edinger-Westphal nucleus. Representative images of the Edinger-Westphal nucleus (EW) from two different biological replicates. No evidence of spatial clustering and uniform distribution of co-infected cells. Scale bars: 100 µm.

**Supplementary Figure S9.**
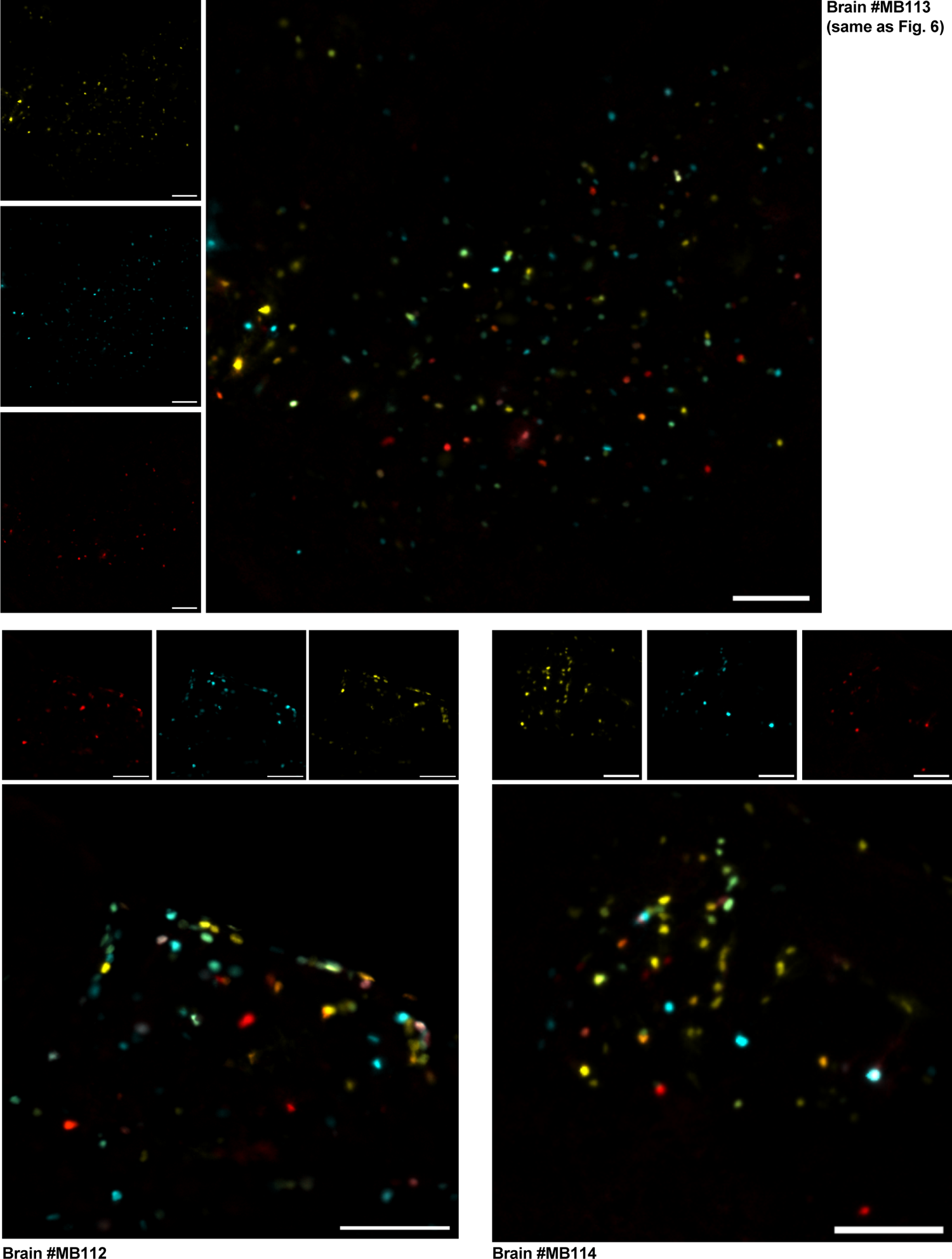
High co-infection levels in the trigeminal nerve nuclei. Representative images of the trigeminal nerve nuclei (TGN) from three different biological replicates. No evidence of spatial clustering and uniform distribution of co-infected cells. Scale bars: 100 µm.

**Supplementary Figure S10.**
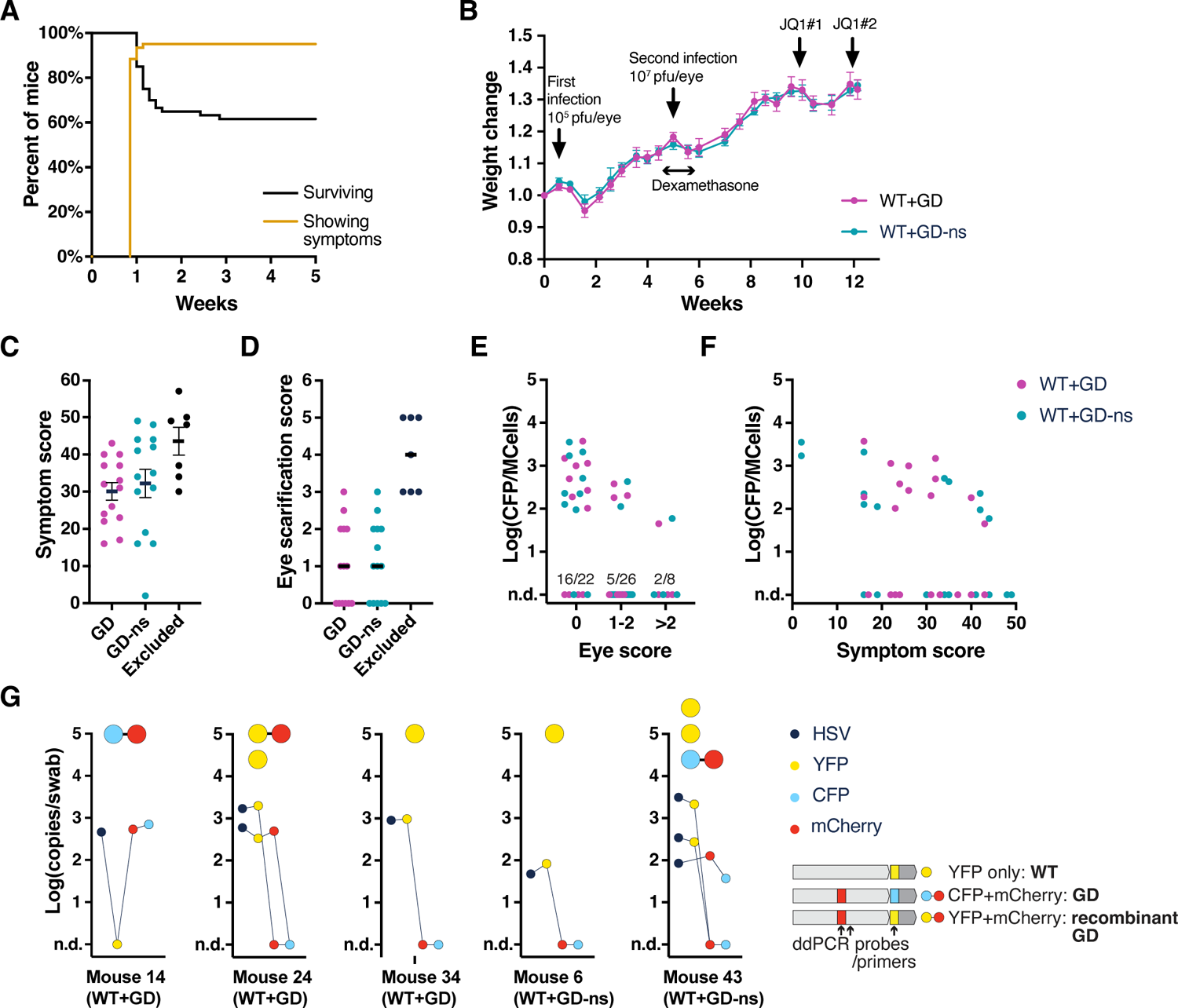
Latent infection in Swiss-Webter mice **A**. Proportion of mice surviving and showing symptoms after primary infection of Swiss-Webster mice with HSV1-WT. n=60 **B**. Weight changes throughout the experiment. **C.** Cumulative symptom score measured during primary infection. Data show means and SEM. **D.** Final eye scarification score at the end of the primary infection. Mice were separated into equivalent groups before superinfection with GD or GD-ns. **E-F**. Final titer of GD/GD-ns in the TG as a function of the eye and symptom scores. **G**. Swab genotyping by duplex ddPCR. Swabs expressing YFP only are wild-type. Swabs expressing CFP and mCherry represent the original GD/GD-ns. Swabs expressing YFP and mCherry are recombinants. Titers are expressed in log-transformed copies per swab, or per million cells after normalization with mouse *RPP30*. n.d.: non-detected.

**Supplementary Figure S11.**
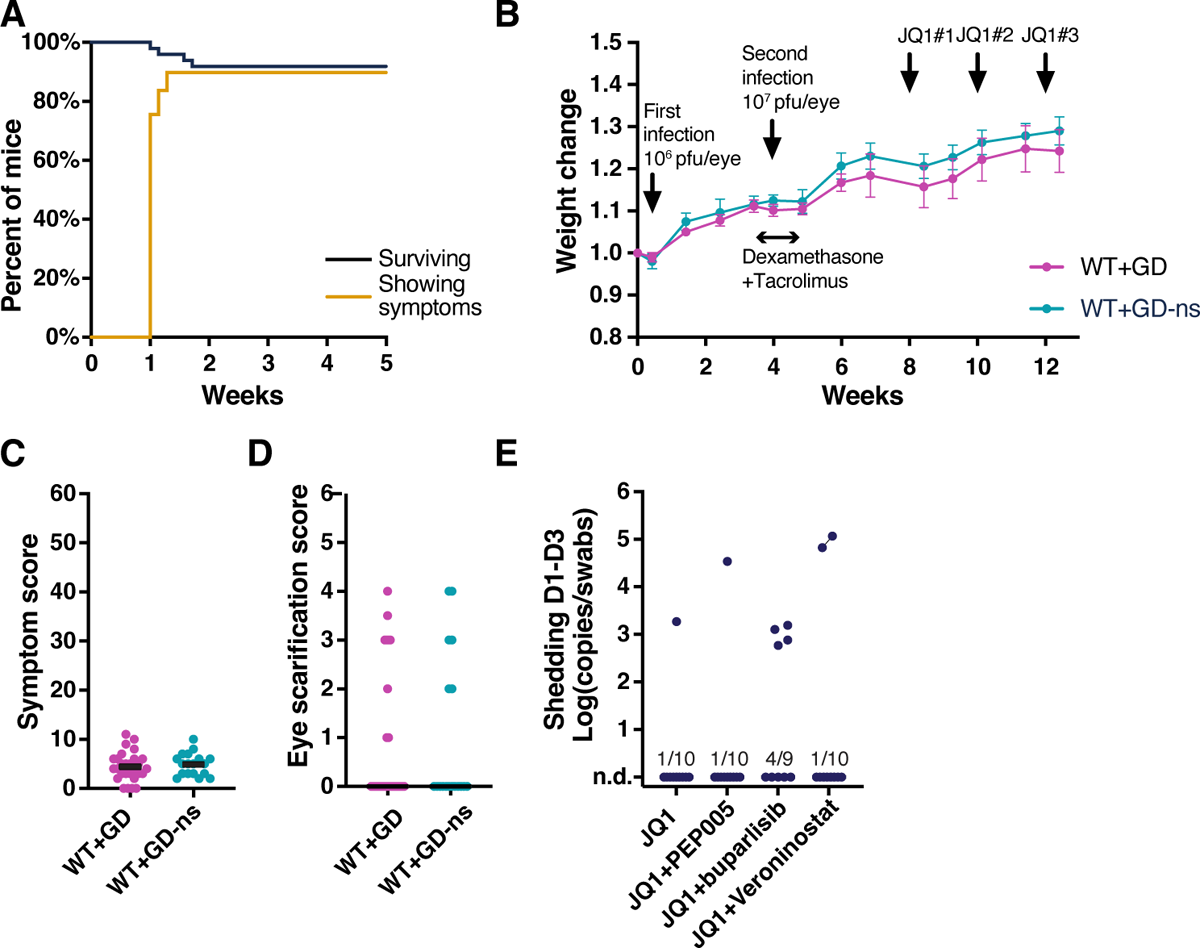
Latent infection in C57Bl/6 mice **A**. Proportion of mice surviving and showing symptoms after primary infection of C57Bl/6 mice with HSV1-WT. n=50 **B**. Weight changes throughout the experiment. C. Cumulative symptom score measured during primary infection. **D.** Final eye scarification score at the end of the primary infection. **E.** Test of different drug combinations to improve reactivation. Based on previous studies, we tested if the small molecules PEP005, Buparlisib or Veroninostat could improve reactivation rates compared to JQ1 alone. In particular, Buparlisib is a brain permeable phosphoinositide 3-kinase (PI3K) inhibitor and PI3K inhibition can reactivate HSV-1 in vitro (34, 35). In an experiment independent from the rest of the study, Swiss-Webster mice latently infected with HSV-1 were treated with JQ1 alone or in combination. Eye swabs collected on days 1-3 were screened by qPCR. Results were not statistically significant but suggested that Buparlisib could improve reactivation rates. Titers are expressed in log-transformed copies per swab. n.d.: non-detected.

**Supplementary Figure S12.**
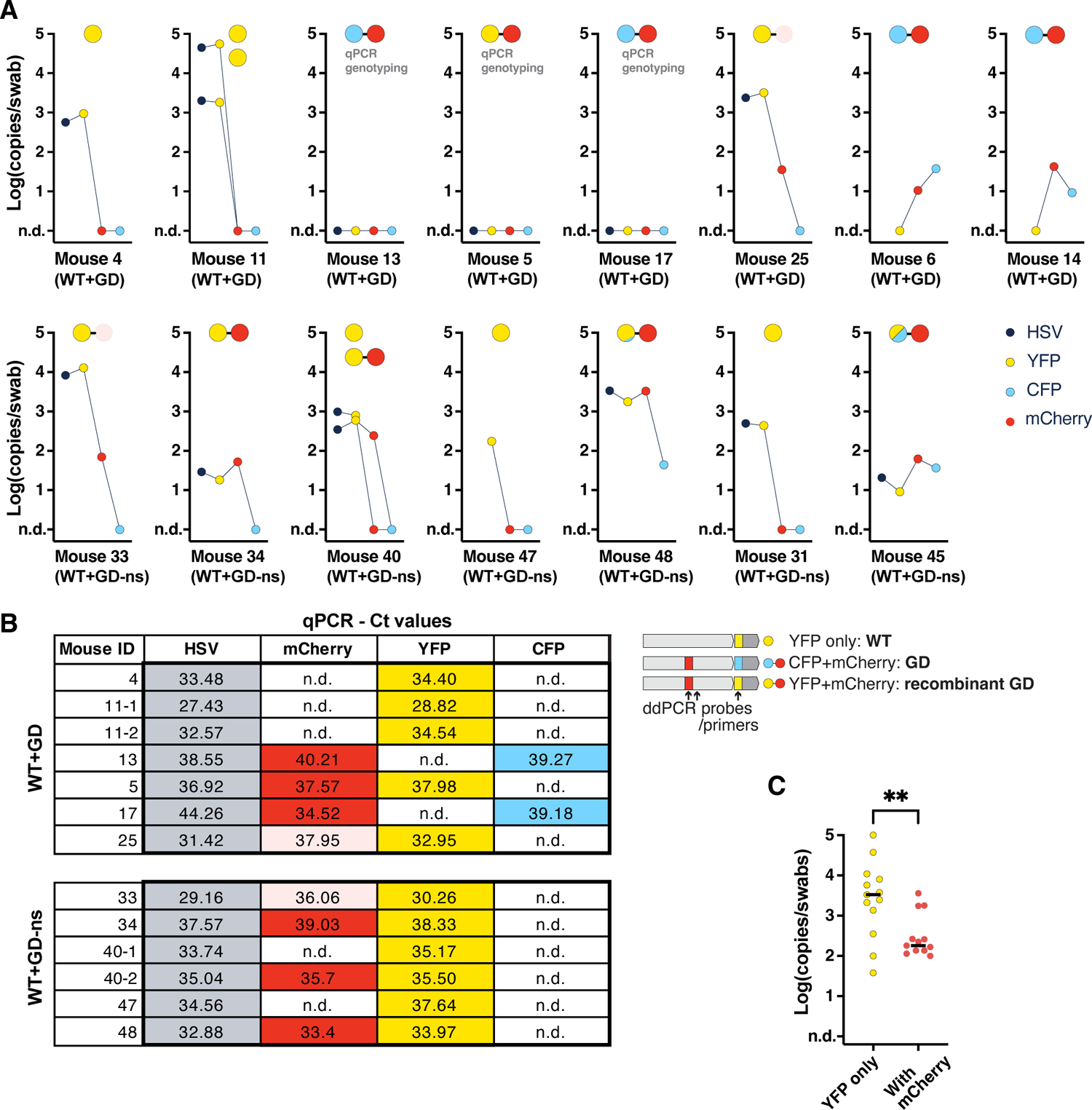
Swab genotyping **A**. Swab genotyping by duplex ddPCR. Swabs expressing YFP only are WT. Swabs expressing CFP and mCherry represent the original GD/GD-ns. Swabs expressing YFP and mCherry are recombinant gene drive viruses. Titers are expressed in log-transformed copies per swab. **B**. Genotype confirmation by duplex qPCR. The table shows CT values. **C.** Swabs genotyped as wild-type (expressing YFP only) had significantly higher titers than swabs genotyped as gene drive (expressing mCherry with either YFP or CFP). Swab qPCR titers from the two experiments in Fig. 7 and 8 were used. Welch’s t-test on log-transformed data, p=0.0079. n=13 for YFP-only, n=12 for mCherry-expressing.

**Supplementary Figure S13.**
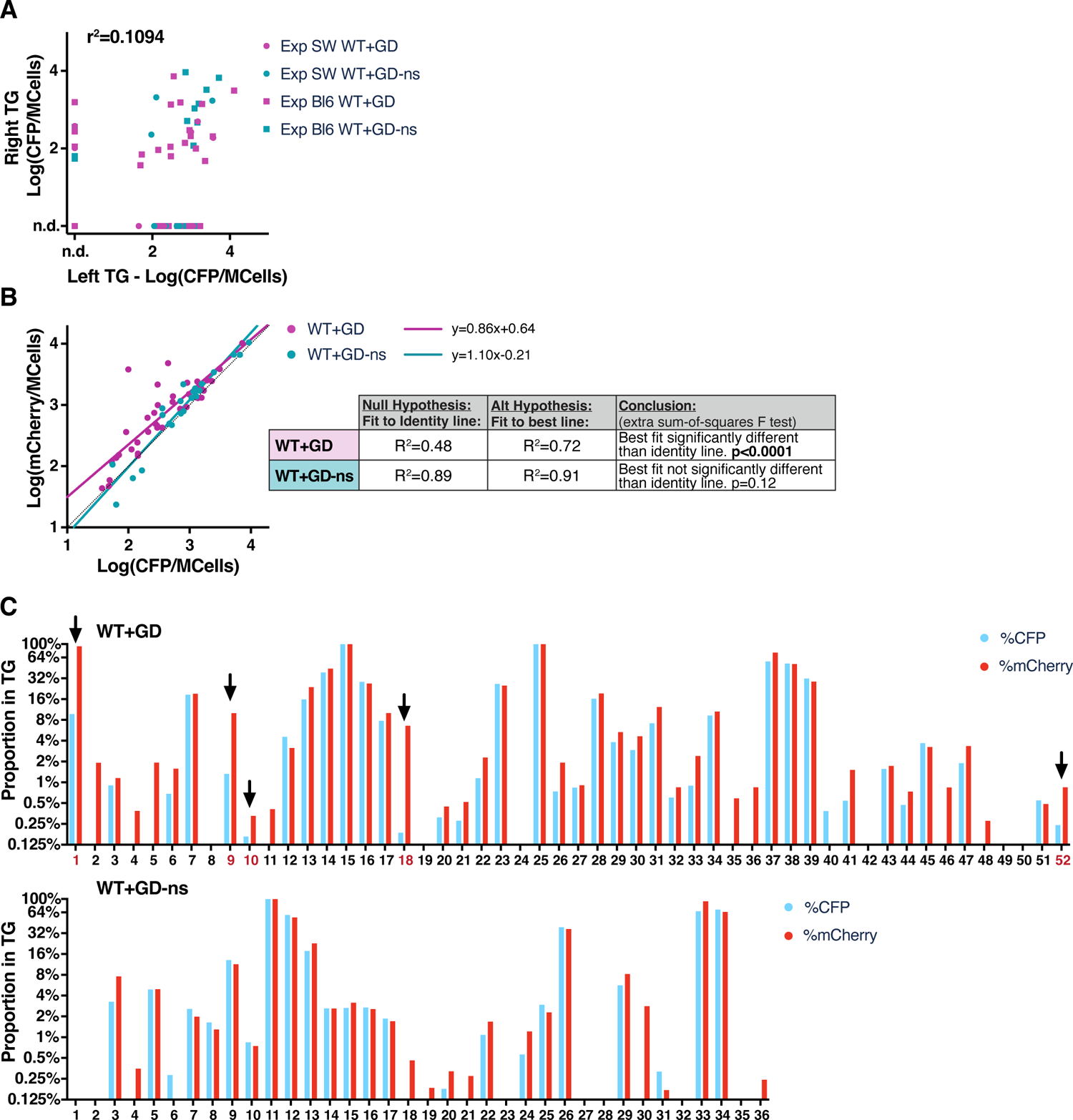
Enrichment of mCherry over CFP in the latent reservoir. **A**. No correlation was observed between CFP titers in the left and right TG, using data from Fig. 7 and 8 using data from Swiss-Webster (SW) and C57Bl/6 (Bl6) mice from Fig. 7 and 8. Pearson correlation coefficient r^2^=0.1094. n=72. **B**. Statistical analysis related to Fig. 8F, showing enrichment of mCherry over CFP in the latent reservoir. Data was fitted either to the best possible line or to the identity line, and the fits were compared using the extra sum-of-squares F test. **C**. Same data as Fig. 8E, showing the proportion of CFP and mCherry in the TG, for each replicate. In the WT+GD-ns control, the proportion of CFP and mCherry are almost equal. In the WT+GD experiment, the proportion of mCherry is consistently higher (70% on average). Black arrows highlight extreme examples with five- to ten-fold enrichment of mCherry over CFP.

